# Chimeric Inheritance and Crown-Group Acquisitions of Carbon Fixation Genes within Chlorobiales

**DOI:** 10.1101/2022.02.08.479544

**Authors:** Madeline M. Paoletti, Gregory P. Fournier

**Affiliations:** Wellesley College; Department of Earth, Atmospheric, & Planetary Sciences, Massachusetts Institute of Technology

## Abstract

The geological record of microbial metabolisms and ecologies primarily consists of stable isotope fractionations and the diagenetic products of biogenic lipids. Carotenoid lipid biomarkers are particularly useful proxies for reconstructing this record, providing information on microbial phototroph primary productivity, redox couples, and oxygenation. The biomarkers okenane, chlorobactane, and isorenieratene are generally considered to be evidence of anoxygenic phototrophs, and provide a record that extends to ∼1.64 Ga. The utility of the carotenoid biomarker record may be enhanced by examining the carbon isotopic ratios in these products, which are diagnostic for specific pathways of biological carbon fixation found today within different microbial groups. However, this joint inference assumes that microbes have conserved these pathways across the duration of the preserved biomarker record. Testing this hypothesis, we performed phylogenetic analyses of the enzymes constituting the reductive tricarboxylic acid (rTCA) cycle in Chlorobiales, the group of anoxygenic phototrophic bacteria usually implicated in the deposition of chlorobactane and isorenieretane. We find phylogenetically incongruent patterns of inheritance across all enzymes, indicative of horizontal gene transfers to both stem and crown Chlorobiales from multiple potential donor lineages. This indicates that a complete rTCA cycle was independently acquired at least twice within Chlorobiales and was not present in the last common ancestor. When combined with recent molecular clock analyses, these results predict that the Mesoproterzoic lipid biomarker record diagnostic for Chlorobiales should not preserve isotopic fractionations indicative of a full rTCA cycle. Furthermore, we conclude that coupling isotopic and biomarker records is insufficient for reliably reconstructing microbial paleoecologies in the absence of a complementary and consistent phylogenomic narrative.

## Introduction

Lipid biomarkers are geochemically stable molecular remnants of organisms, frequently used to trace paleoenvironmental proxies, or as “fossils” indicating the presence of specific groups of organisms^1–3^. Phototrophic bacteria, including oxygenic cyanobacteria and anoxygenic green sulfur bacteria (GSB, order Chlorobiales) and purple sulfur bacteria (PSB, order Chromatiales), have especially significant biomarker records in the form of carotenoid pigment derivatives. These can indicate periods of major planetary change, such as great oceanic anoxic events^4^.

The first appearance of sulfur bacteria in the biomarker record is from the 1.64 Ga Barney Creek Formation, which contains carotenoids including okenane, chlorobactane, and isorenieratane^5,6^. This implies sulfur bacteria were a substantial part of aquatic microbial ecologies by this time, consistent with the presence of a stratified ocean with widespread euxinic conditions^7^. However, the genes encoding the biosynthesis of aromatic carotenoids and their derivatives have also been identified in cyanobacteria, which occupy more oxygenated water columns^8–10^. Furthermore, new geochemical arguments have suggested a cyanobacterial origin for the fossil biomarkers found in Proterozoic depositional settings^11^. This brings the presence of GSB at 1.64 Ga into question, as well as the practice of using these biomarkers as a calibration for dating Chlorobiales in molecular clock analyses^12,13^.

A distinct, comparable source of information for tracing microbial autotrophy can be found within the geochemical record of carbon fixation, which can further inform reconstructions of microbial paleoecology. There are six known modern biological pathways used by organisms to fix CO_2_. Most fractionate carbon differently, variably offsetting carbon isotope (δ^13^C) values of inorganic and organic carbon in the isotopic record^14,15^. Through this bulk signaling of δ^13^C sources, a cycle’s input can be mathematically inferred by isotopic mass balance of its components^16^.

Of the known carbon fixation pathways, the Calvin-Benson-Bassham (CBB) cycle and the rTCA cycle are the most prevalent. The taxonomic distributions of these pathways are polyphyletic, consistent with deep evolutionary histories of horizontal gene transfer (HGT)^16,17^. The CBB cycle is found in photoautotrophic eukaryotes, cyanobacteria, and Chromaticeae, and fixes three molecules of CO_2_ for the synthesis of one 3-phoshoglycerate molecule using ribulose-1,5-bisphosphate carboxylase/oxygenase (RuBisCo). The rTCA cycle is present in a diverse group of autotrophic microbes, including Chlorobiaceae, sulfate-reducing bacteria, hyperthermophilic bacteria, and Crenarchaeota^18,19^. The complete cycle fixes three molecules of CO_2_ for the synthesis of one 3-carbon pyruvate molecule through 9 steps that also produce several metabolic intermediates^20–22^ (Figure 1). Except for ATP-citrate lyase, which drives the reductive cycling^23^, and pyruvate: ferredoxin oxioreductase and 2-oxoglutarate:ferredoxin oxioreductase (OGOR) needed to reduce CO_2_, these enzymes are also used within the oxidative TCA cycle, which has been shown to operate within GSB under some growth conditions^24^. However, a complete oxidative TCA cycle is not present in obligate anaerobes, as aerobic respiratory chains are presumably necessary for the efficient re-oxidation of NADH^25^.

**Figure 1.**
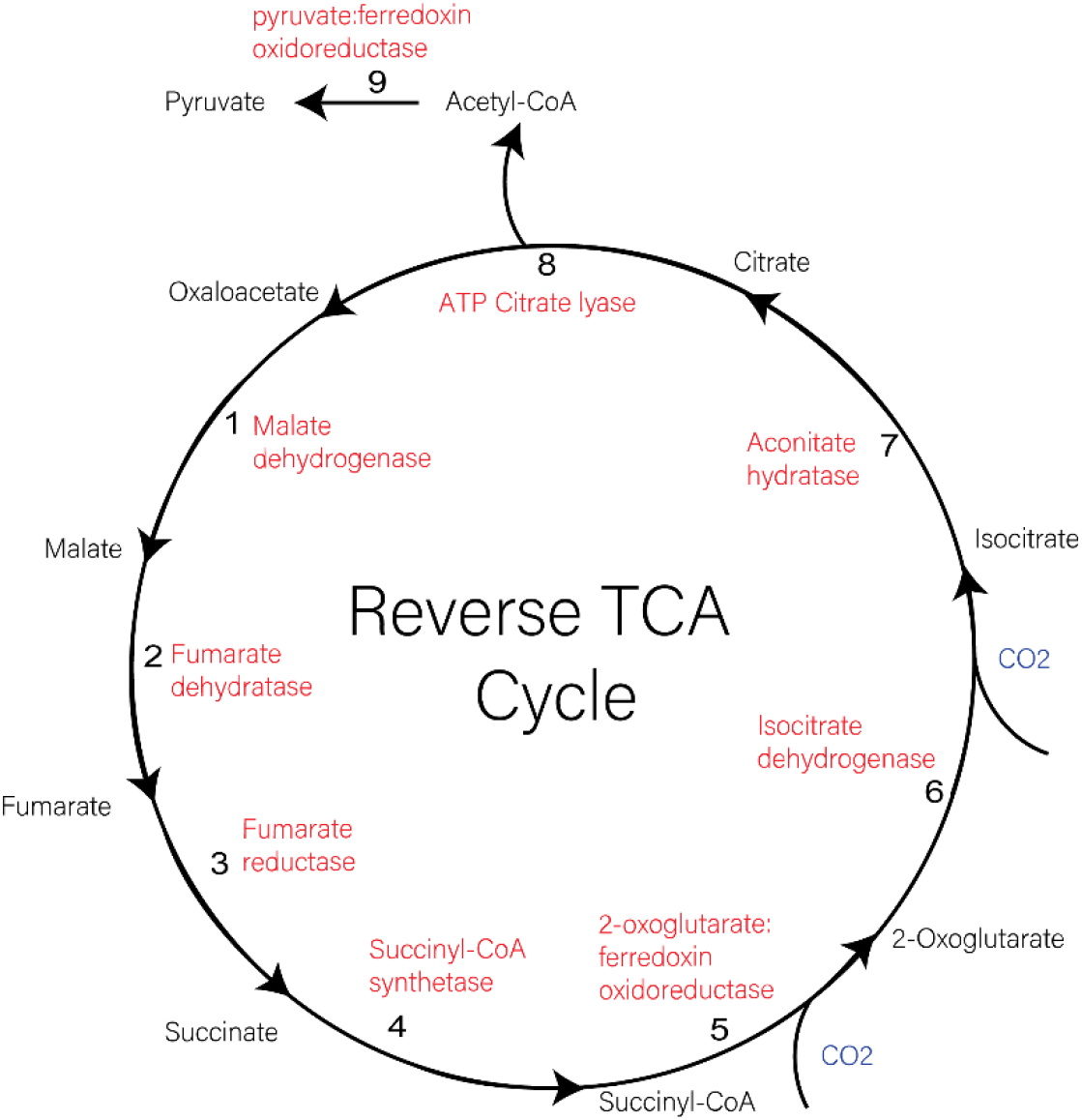
Enzymatic map of the rTCA cycle. Enzymes catalyzing each reaction (red), inputs of carbon dioxide (blue) and intermediate compounds (black) are shown. Steps are numbered in the direction of reactions, starting with the conversion of oxaloacetate to malate via malate dehydrogenase.

Recently, an alternative rTCA cycle has been characterized that uses two different enzymes to convert citrate into citryl-CoA, instead of ATP citrate lyase^26^. This was initially observed in *Hydrogenobacter thermophilus* but is also present in other members of Aquificeae^27^. Anaerobes with partial TCA cycles also use α-ketoglutarate:ferredoxin oxioreductase, and fumarate reductase instead of succinate dehydrogenase, similar to the rTCA cycle^25,28^, and pyruvate: ferredoxin oxioreductase is used in some carbohydrate fermentation pathways^29^. Thus, the phyletic distribution of genes found in the rTCA cycle is complex, and potentially indicative of several different metabolic processes other than autotrophy.

The order Chlorobiales consists of a single family (Chlorobiaceae) and contains five genera (Chlorobium, Chloroherpeton, Pelodityon, Ancalochloris, and Prosthecochloris) and a group of unclassified thermophiles^30^. The 16S rRNA phylogenies of this group recover close relationships to Ignavibacterales (within the phylum Chlorobi) and the Bacteriodetes, which are often grouped together along with Firmicutes in a CFB superphylum^31,32^. Metagenomic analyses and culturing of novel strains has revealed more aerobic and photoheterotrophic members of Chlorobi lacking enzymes for autotrophic carbon-fixation^33–36^. Some of these uncultured genome species, such as *Candidatus Thermochlorobacter aerophylum*^33,34^, *Chlorobium* sp. 445 and *Chlorobium* sp. GBChlB^35^ group as a distinct clade within Chlorobiaceae. Individual gene phylogenies incongruent with this species-tree history could potentially infer the transfer history to both stem and crown Chlorobiales.

If the constituent enzymes of the rTCA cycle were acquired within crown Chlorobiales, rather than the stem lineage, then ancestral autotrophy cannot be inferred for this order. Subsequently, diagnostic lipid biomarkers more ancient than the inferred age of crown Chlorobiales should not show isotopic fractionations indicative of rTCA cycle carbon fixation. Conversely, a phylogenetic signal showing clear acquisition of all rTCA enyzmes within stem Chlorobiales would be supportive of a much older history of autotrophy within this group and would predict isotopic fractionations of even the oldest preserved biomarkers to be similar to those of modern autotrophic Chlorobiaceae. To investigate this, we performed phylogenetic analyses of rTCA cycle enzymes in Chlorobiales to map their evolutionary histories. Overall, our investigations reveal a mixed history of vertically inherited and horizontally transferred rTCA cycle genes across multiple groups of Chlorobiales (Table 1). These complex histories support a hypothesis of patchwork enzymatic inheritances from multiple and diverse HGT donors, implying the complete rTCA cycle, and therefore autotrophy, was not present in the ancestral stem group of Chlorobiales.

**Table 1:** Summary of evolutionary histories of carbon fixation genes in Chlorobiaceae. Italicized species names refer to taxon source of query sequence for respective groupings.

| rTCA Step | Enzyme Name | Inferred history | Sibling Groups | Outgroup Carbon Metabolism |
| --- | --- | --- | --- | --- |
| 1 | <u>malate dehydrogenase</u> | Chlorobiales: stem HGT | *Rhodothermus marinus* | Mixed |
| 2 | <u>fumarate hydratase</u> | Chlorobiales: stem HGT or vertical inheritance<br>*C.thalissium*: orthologous displacement | Bacteroidetes<br>Ignavibacterium<br>Plantomycetaeaceae bacterium<br>Oceanospiralles bacterium | Heterotrophs<br>Heterotrophs |
| 3 | <u>fumarate reductase</u> | Chlorobiales: stem HGT | Candidatus<br>Lambdaproteobacteria<br>*Geobacter*<br>Desulfuromonadales bacterium | Mixed |
| 4 | <u>succinyl-CoA synthetase</u> | Chlorobiales: vertical inheritance | Bacteroidetes<br>Ignavibacterium | Heterotrophs |
| 5 | <u>2oxoglutarate:ferredoxin oxidoreductase</u> | Chlorobiales: stem HGT from Bacteroidetes | Balneolaceae<br>Bacteroidetes | Heterotrophs |
| 6 | <u>isocitrate dehydrogenase</u> | Chlorobiales: HGT<br>*C.thalissium*/Photoheterotrophs: HGT | Deltaproteobacteria<br>Nitrospira/Nitrospinae<br>Armatimonadetes<br>Caldithrix | Mixed<br>Mixed |
| 7 | <u>aconitate hydratase</u> | Other Chlorobiales: HGT<br>Chloroherpeton: HGT<br>Thermochlorobacter: HGT | Campylobacterales<br>Ignavibacteria<br>Spirochaetes | Autotrophs<br>Heterotrophs |
| 8 | <u>ATP citrate lyase</u> | Chlorobiales: stem HGT | Nitrospirae<br>Nitrospinae | Mixed |
| 9 | <u>pyruvate:ferredoxin oxidoreductase</u> | Chlorobiales: HGT | Bacteroidetes<br>Ignavibacteria | Mixed |

## Results

### rTCA cycle protein phylogenies show a complex history of HGT

Maximum-likelihood phylogenies of rTCA pathway proteins within Chlorobiales show surprisingly diverse evolutionary origins (Table 1). Several show monophyly indicative of acquisition within stem Chlorobiales (malate dehydrogenase, pyruvate ferredoxin, ATP citrate lyase, succinyl-CoA synthetase, OGOR, fumarate reductase). Other enzymes show polyphyletic distributions, indicative of multiple HGT acquisitions from different donor lineages (isocitrate dehydrogenase, fumarate hydratase, aconitate hydratase). Several enzymes were subsequently lost within the photoheterotrophic Chlorobiales clade (aconitate hydratase, pyruvate: ferredoxin oxidoreductase, fumarate reductase, ATP-citrate lyase), likely following divergence from the *Chloroherpeton* lineage.

Several general trends can be identified in the placement of these groups within broader protein diversity, based on the outgroup sequences within each tree. There are three scenarios wherein the inheritance of an rTCA cycle protein could be inferred for crown Chlorobiales. First, trees where the outgroups are similar to those of the species tree, such as other Chlorobi (e.g., Ignavibacteria and/or Bacteroidetes) suggest that these proteins were ancestral within the Chlorobiales lineage, and subsequently vertically inherited. Of the rTCA pathway proteins, only succinyl-CoA synthetase appears to have a history consistent with this interpretation. Second, trees where Chlorobiales sequences are on long branches without any closely related outgroups and crown group monophyly is preserved (malate dehydrogenase, pyruvate:ferredoxin, ATP citrate lyase). These enzymes were likely acquired by stem Chlorobiales via HGT, but current taxonomic sampling and/or lack of phylogenetic signal prevent an unambiguous identification of a putative donor lineage. Third, trees showing vertical inheritance within crown Chlorobiales following an HGT from a clear donor group to stem Chlorobiales. Only OGOR preserves this signal. The remaining tree topologies are complicated by HGTs within crown group lineages, so that Chlorobiales is polyphyletic within the tree (fumarate hydratase, isocitrate dehydrogenase, aconitate hydratase). Either these genes were acquired in stem Chlorobiales and a subsequent orthologous displacements occurred, or the original acquisitions within crown Chlorobiales were independent.

### Malate Dehydrogenase

The ML tree for malate dehydrogenase shows a broad taxonomic range of bacteria, with 11 distinct phyla represented in 242 taxa. The tree is consistent with acquisition of malate dehydrogenase in the Chlorobiales stem ancestor, and subsequent vertical inheritance. The sibling groups are diverse, including both autotrophic and heterotrophic representatives of Gemmatimonadetes, Deltaproteobacteria, Acidobacteria, and Bacteroidetes, with poorly supported relationships between most groups (Figure 2A). This indicates a complex history of HGT between phyla, preventing identification of a likely HGT donor to stem Chlorobiales.

**Figure 2.**
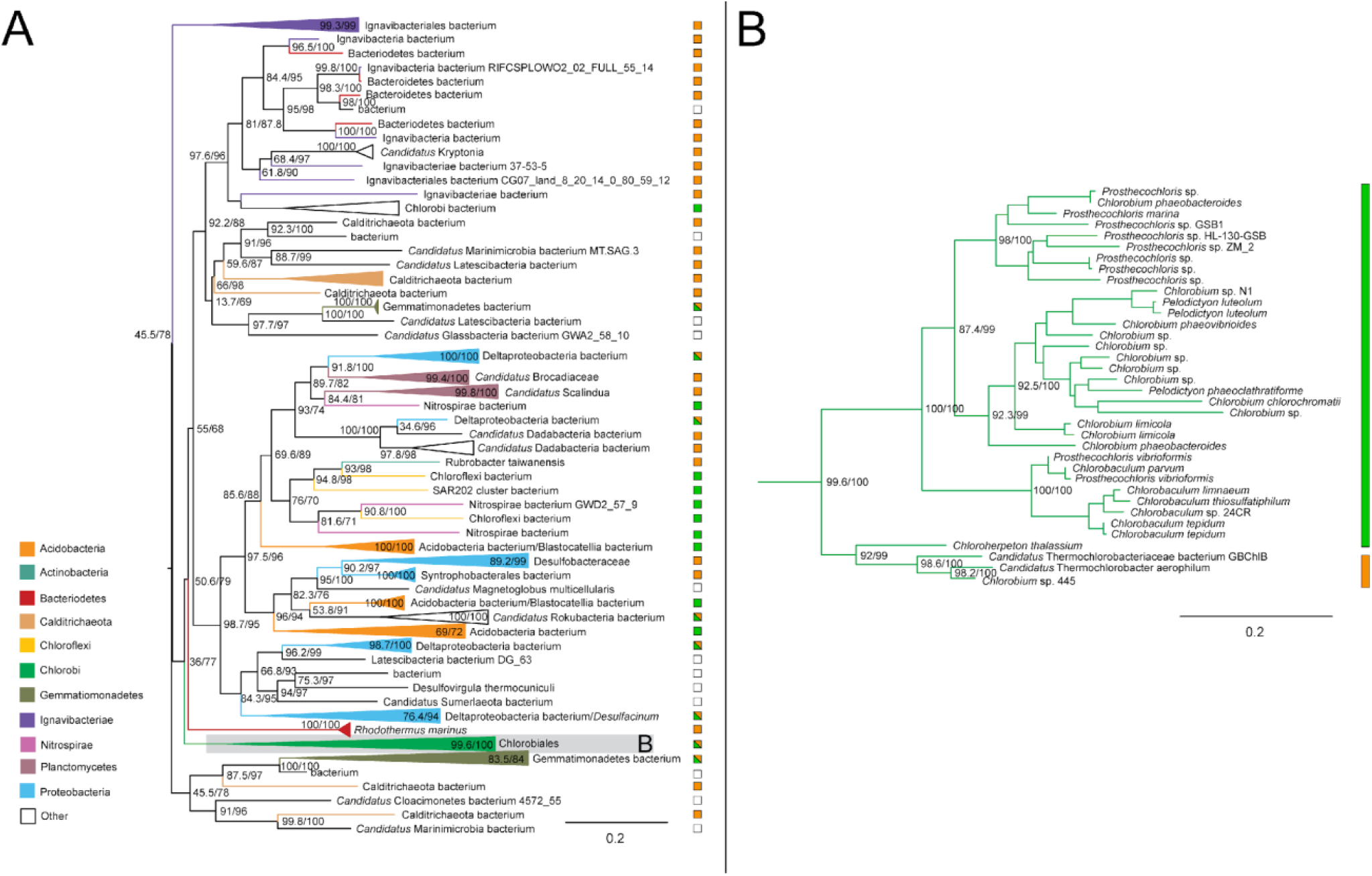
Maximum likelihood (ML) tree of malate dehydrogenase homologs. (A) midpoint-rooted tree with collapsed clades labeled with taxonomic group names. (B) Higher resolution tree showing crown Chlorobiales. Support values indicate approximate likelihood ratio test (aLRT)/ bootstrap (100 replicates). Major bipartitions with bootstrap (BS) support are labeled. The full tree contains 242 taxa. Collapsed clades are labeled with taxonomic group names. Color bars or boxes to the right of the trees indicate autotrophic (green), heterotrophic (orange), or undetermined (white) carbon metabolisms. Clades including multiple carbon metabolisms are indicated with diagonally hatched boxes.

### Fumarate Hydratase

Genes encoding fumarate hydratase do not recover the monophyly of Chlorobiales, with the ortholog from *Chloroherpeton thalissium* more closely related to homologs from other bacterial groups. This is consistent with two possible evolutionary scenarios: (1) ancestral presence and vertical inheritance within GSB (with the exception of *C*.*thalissium*) in which the gene underwent subsequent orthologous displacement; (2) independent acquisition via two independent HGT events following divergence of *C*.*thalissium* from the ancestor lineage of other Chlorobiales. In either scenario, the *C*.*thalissium* copy of the gene was likely acquired from within gammaproteobacteria; some members of Chloroflexi and Acidobacteria appear to have acquired this gene within gammaproteobacteria as well (Figure 3A). *Chlorobium* sp. 445 and *Candidatus Thermochlorobacter*, which usually group with *C*.*thalissium* on other trees, do not for this gene, which further indicates an orthologous displacement for *C*.*thalissium* after its divergence from the photoheterotrophic lineages. The broad and sparse taxonomic distribution of sequences grouping with crown Chlorobiales (Figure 3B) prevents the identification of a likely HGT donor group. or an inference of vertical inheritance in stem Chlorobiales, even though some Ignavibacteria sequences are present.

**Figure 3.**
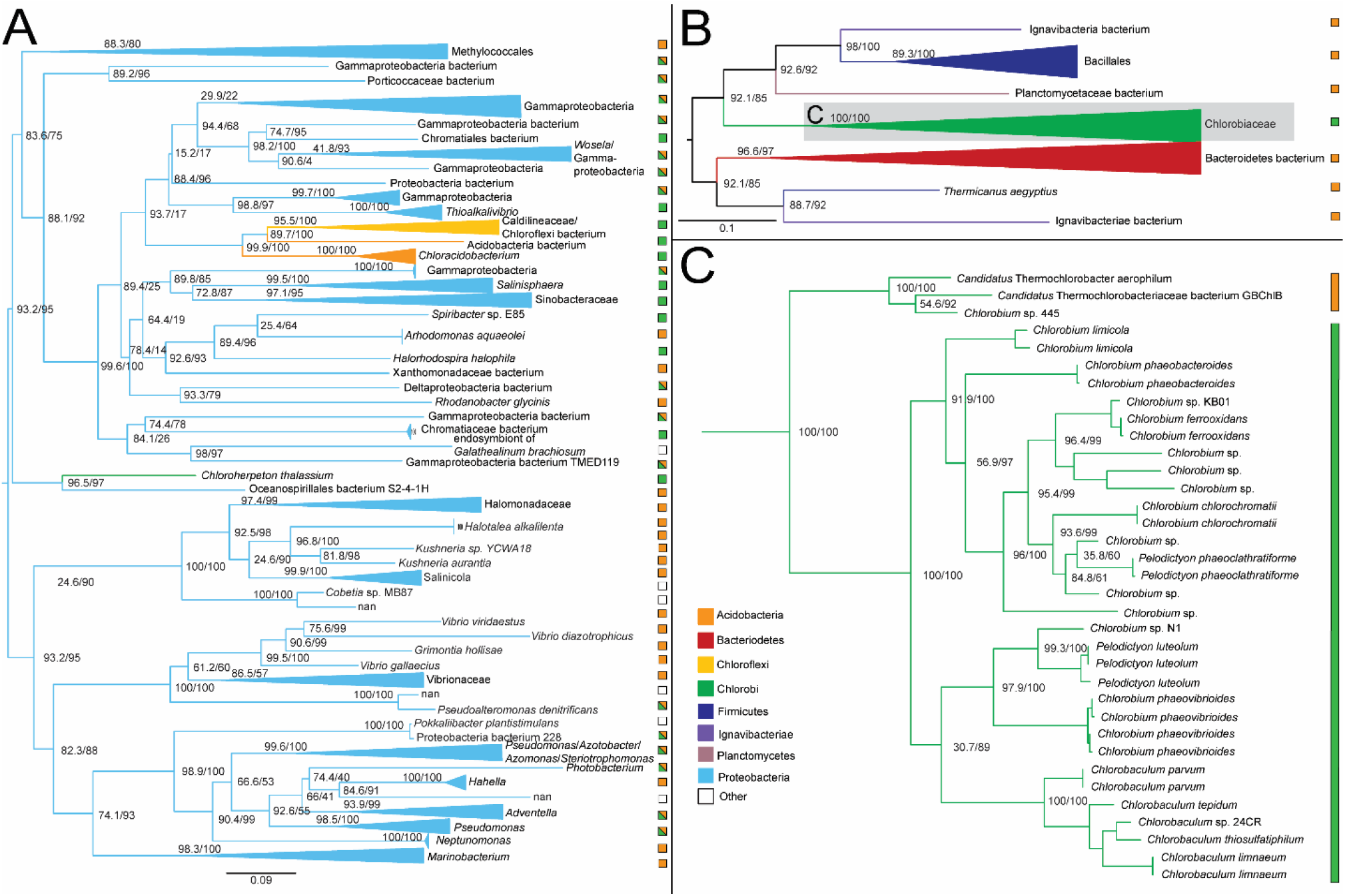
Maximum likelihood (ML) trees of fumarate hydratase homologs. (A-B) Two groups of sequences related to Chlorobiales query genes were identified, shown as midpoint rooted trees with collapsed clades labeled with taxonomic group names. (C) Higher resolution tree of crown Chlorobiales sequences from B. Support values indicate approximate likelihood ratio test (aLRT)/ bootstrap (100 replicates). Major bipartitions with bootstrap (BS) support are labeled. The full trees contain 250 and 242 taxa, respectively. Color bars to the right of the tree indicate autotrophic (green), heterotrophic (orange), or undetermined (white) carbon metabolisms. Clades including multiple carbon metabolisms are indicated with diagonally hatched boxes.

### Fumarate Reductase

The ML tree of fumarate reductase homologs recovers the monophyly of Chlorobiales sequences, closely related to sequences from Candidatus Lambdaproteobacteria, *Geobacter*, and Desulfuromonadales bacterium (Figure 4). These are distantly related by a long branch to other bacterial sequences, including members of Firmicutes, Actinobacteria, and other Proteobacteria, indicative of an HGT into stem Chlorobiales, although the donor group cannot be discerned. The short branch separating the Chlorobiales and proteobacterial sequences indicates a history of relatively recent HGT involving stem Chlorobiales, although the absence of a polarizing outgroup prevents the direction of HGT from being established. A stem Chlorobiales lineage could have been the primary HGT donor to this group of Proteobacteria, or vice versa. Both groups could, alternatively, have been HGT recipients from an unsampled or extinct lineage, potentially explaining the long empty branch preceding this grouping.

**Figure 4.**
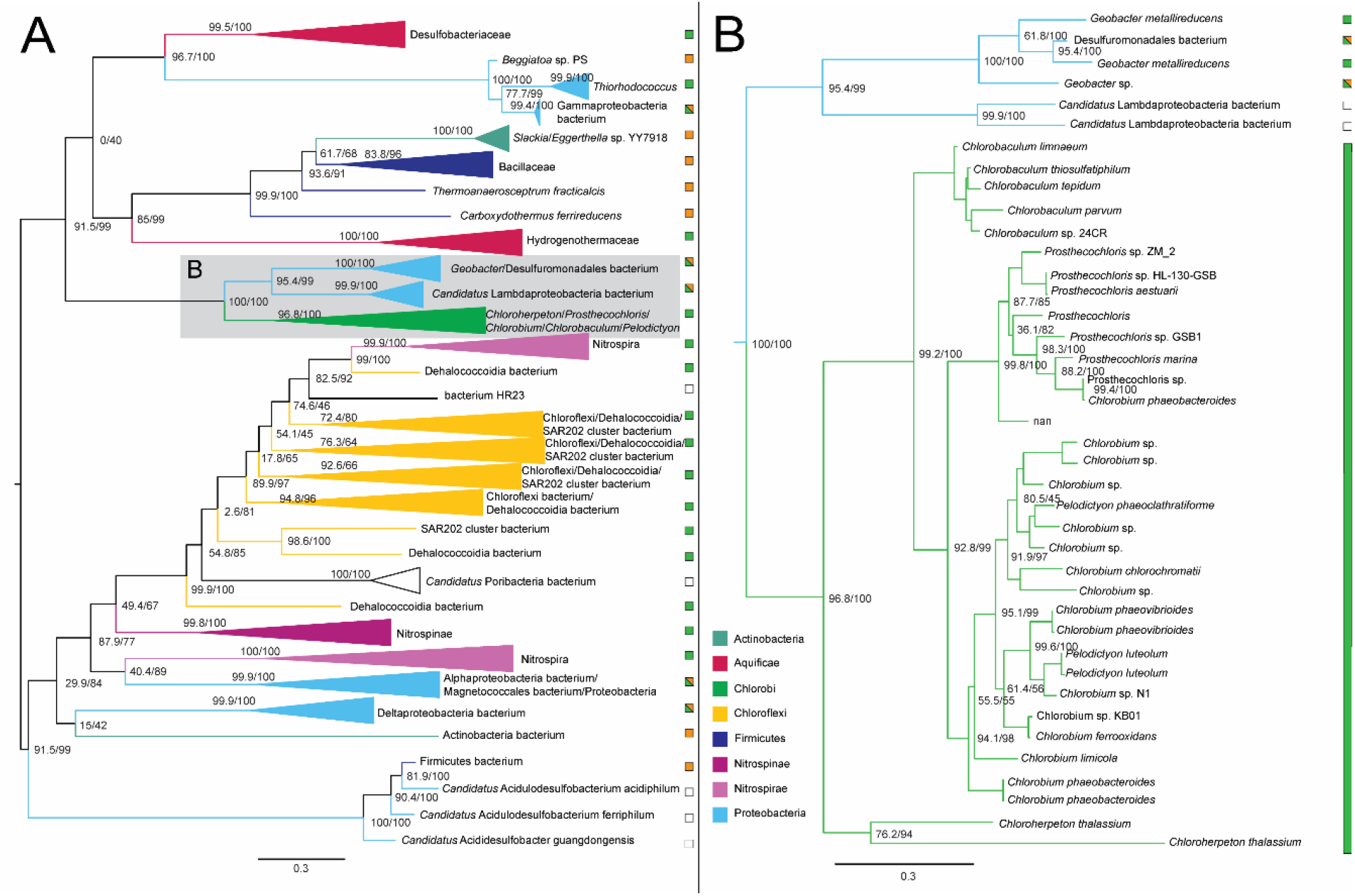
Maximum likelihood (ML) tree of fumarate reductase homologs. (A) midpoint-rooted tree with collapsed clades labeled with taxonomic group names. (B) Higher resolution tree of crown Chlorobiales and closely related proteobacterial sequences. Support values indicate approximate likelihood ratio test (aLRT)/ bootstrap (100 replicates). Major bipartitions with bootstrap (BS) support are labeled. The full tree contains 248 taxa. Color bars to the right of the tree indicate autotrophic (green), heterotrophic (orange), or undetermined (white) carbon metabolisms. Clades including multiple carbon metabolisms are indicated with diagonally hatched boxes.

**Figure 5.**
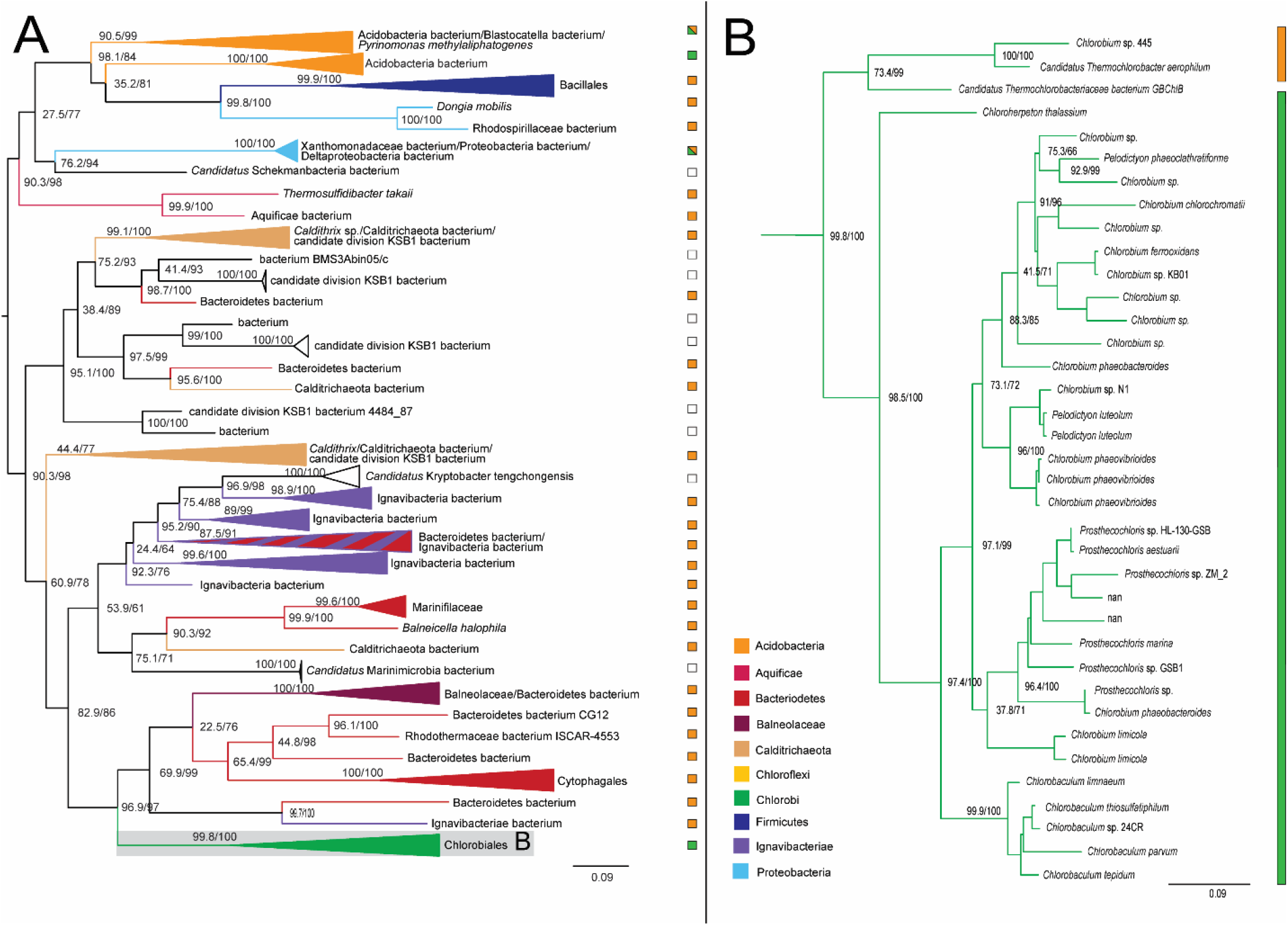
Maximum likelihood tree of succinyl-CoA synthetase homologs. (A) midpoint-rooted tree with collapsed clades labeled with taxonomic group names. (B) Higher resolution tree of crown Chlorobiales. Support values indicate approximate likelihood ratio test (aLRT)/ bootstrap (100 replicates). Major bipartitions with bootstrap (BS) support are labeled. The full tree contains 248 taxa. Color bars to the right of the tree indicate autotrophic (green), heterotrophic (orange), or undetermined (white) carbon metabolisms. Clades including multiple carbon metabolisms are indicated with diagonally hatched boxes.

**Figure 6.**
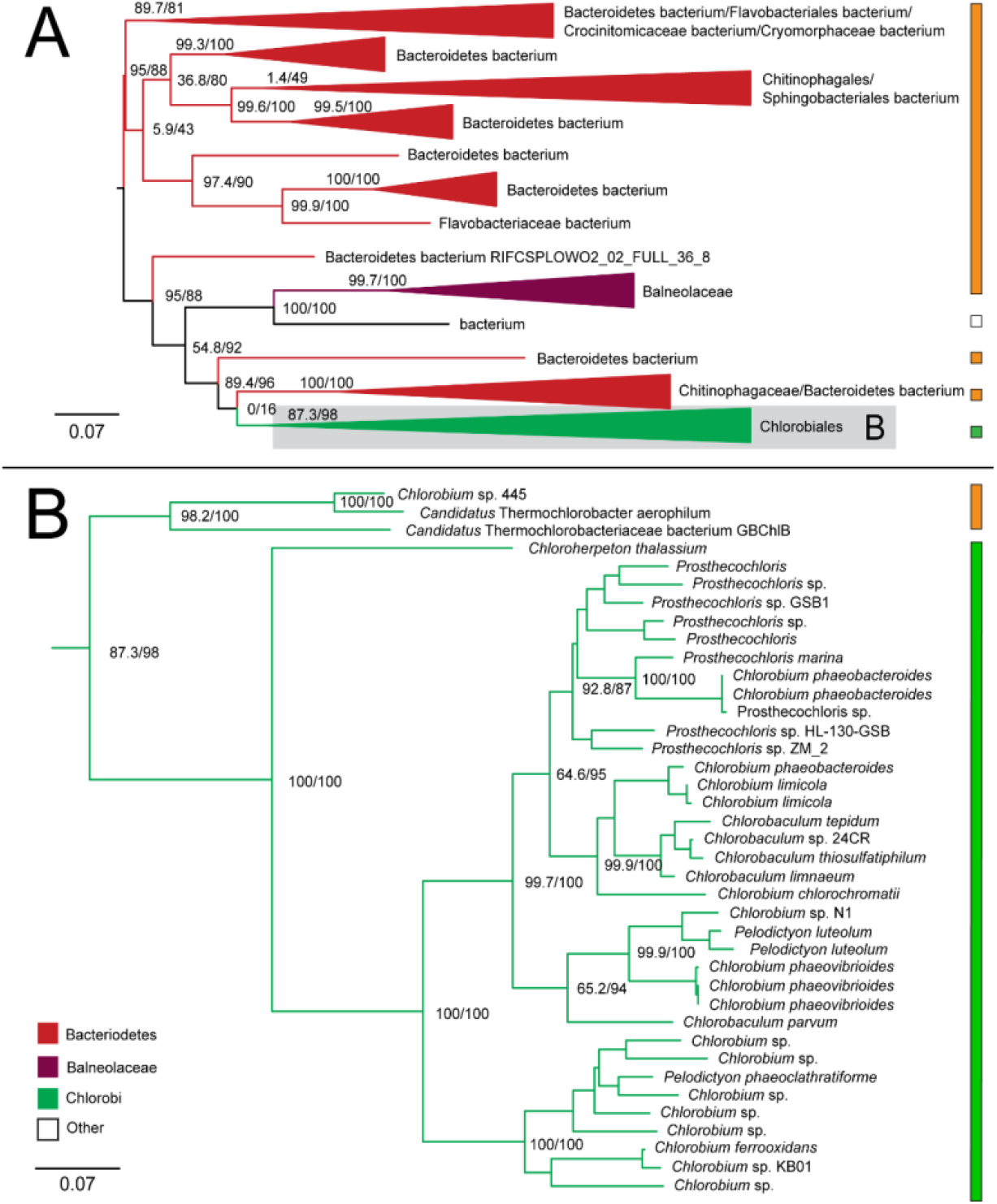
Maximum likelihood (ML) tree of 2-oxoglutarate: ferredoxin oxidoreductase homologs. (A) midpoint-rooted tree with collapsed clades labeled with taxonomic group names. (B) Higher resolution tree of crown Chlorobiales. Support values indicate approximate likelihood ratio test (aLRT)/ bootstrap (100 replicates). Major branches with bootstrap (BS) support are labeled with respective values. Tree A contains 243 taxa. Color bars to the right of the tree indicate autotrophic (green), heterotrophic (orange), or undetermined (white) carbon metabolisms. Clades including multiple carbon metabolisms are indicated with diagonally hatched boxes.

### Succinyl-CoA synthetase

The ML tree for succinyl-CoA synthetase recovers the monophyly of Chlorobiales with a well-supported sibling group including Cytophagales, Marinifilaceae, other Bacteroidetes, and Ignavibacteria. While this outgroup contains similar taxa to the species tree outgroup of Chlorobiales, the phylogeny shows a mixing of Ignavibacteriales and Bacteroidetes groups that prevents inference of vertical inheritance in the Chlorobiales stem ancestor, or identification of an HGT donor group. Photoheterotrophic Chlorobiales have also retained this gene, indicating that its presence is not strictly indicative of a functional rTCA cycle. Succinyl-CoA may be involved in any number of additional metabolic functions, including heme synthesis, ketone metabolism, and full or partial oxidative TCA cycles^37^. More distantly related outgroups on the tree also show extensive HGT between bacterial groups, including Calditrichaeota, Firmicutes, Acidobacteria, and Proteobacteria.

### 2-oxoglutarate: ferredoxin oxidoreductase

The ML tree for 2-oxoglutarate: ferredoxin oxidoreductase alpha subunit recovers the monophyly of Chlorobiales, grouping within a taxonomically broad range of Bacteroidetes. The placement of the specific orders Chitinophaga, Flavobacteriales, Sphingobacteriales, and Balneolaceaes within the tree suggests similarity to published species trees of Bacteroidetes^32^, and is further indicative of a likely HGT from within Bacteroidetes. The ML tree does not recover the sibling grouping of *C*.*thalassium* and thermophilic photoheterotrophic lineages, suggesting a possible misrooting of Chlorobiales in this tree.

### Isocitrate Dehydrogenase

The isocitrate dehydrogenase phylogeny recovers a polyphyletic placement of sequences within Chlorobiales, with *Chlorobium, Pelodictyon*, and *Chlorobaculum* genera and *Prosthecochloris* genera forming two distinct groups within one set of identified homologs (Figure 7A) and a group containing *C*.*thalassium, Thermochlorobacter*, and C*hlorobium* sp.445 within another set of identified homologs (Figure 7B), providing clear evidence of multiple HGT events for this gene family. Specific HGT donor groups to Chlorobiales cannot be clearly identified from these trees. The placement of closely related Chlorobiales genera in Fig. 7A is consistent with vertical inheritance in this group, with subsequent multiple HGTs to other bacterial groups, including Deltaproteobacteria, Desulfobulbaceae bacterium, and Marinifilaceae. These distinct gene histories suggest that the clades within Chlorobiales shown in Fig. 7C and Fig. 7D acquired isocitrate lyase independently after they diverged.

**Figure 7.**
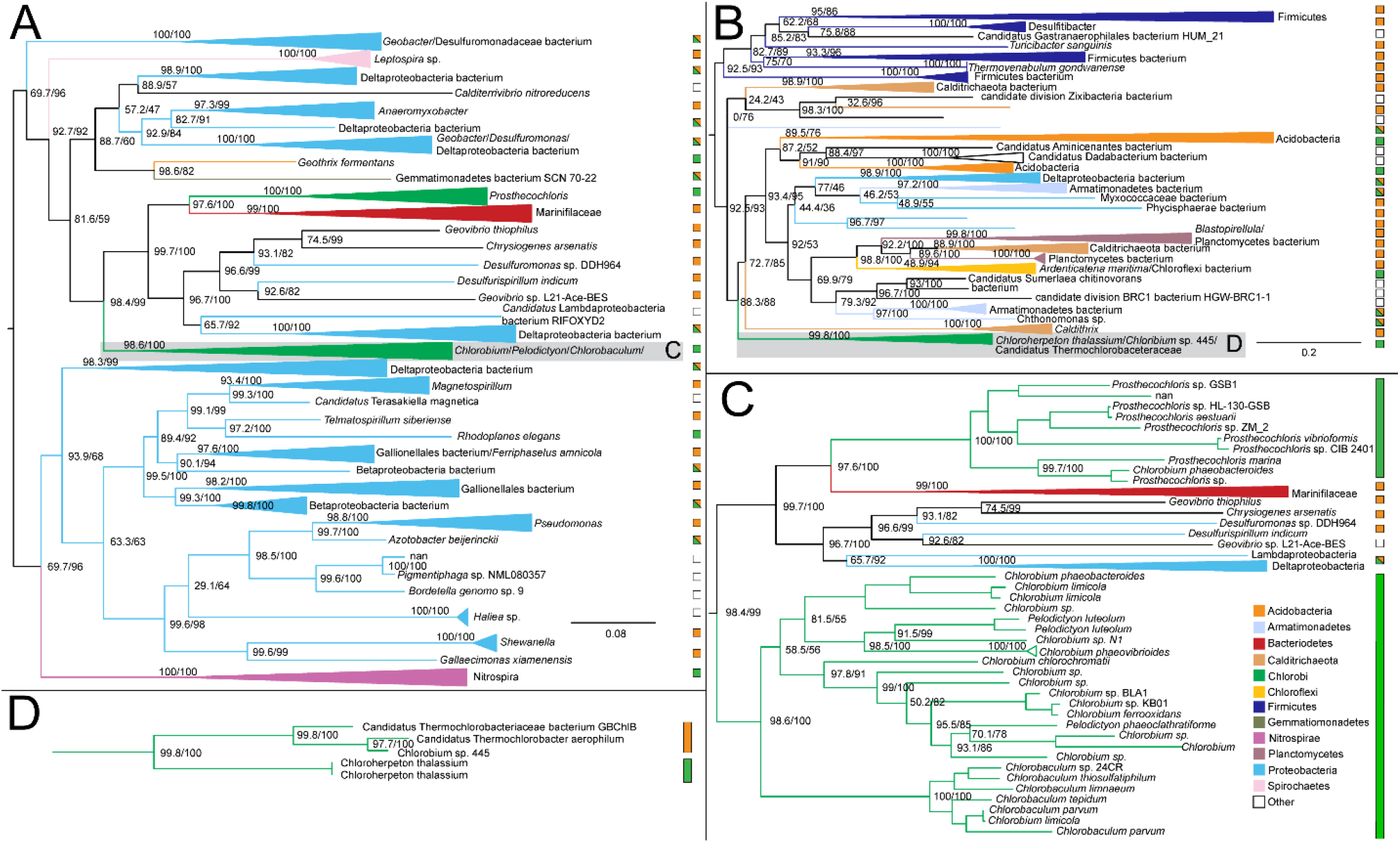
Maximum likelihood (ML) trees of isocitrate dehydrogenase homologs. Two groups of sequences related to Chlorobiales query sequences were identified, shown as midpoint rooted trees with collapsed clades labeled with taxonomic group names (A-B). (C-D) Higher resolution trees of crown Chlorobiales groups. Support values indicate approximate likelihood ratio test (aLRT)/ bootstrap (100 replicates). Major bipartitions with bootstrap (BS) support are labeled. A and B trees contain 250 and 232 taxa, respectively. Color bars to the right of the tree indicate autotrophic (green), heterotrophic (orange), or undetermined (white) carbon metabolisms. Clades including multiple carbon metabolisms are indicated with diagonally hatched boxes.

### Aconitate hydratase

Aconitate hydratase sequences within Chlorobiales are polyphyletic with *C. thalassium, Thermochlorobacter*, and remaining taxa each grouping with different sets of bacterial sequences. The main group (Figure 8C) places within Epsilonproteobacteria, suggesting this is the HGT donor group to the major clade of crown Chlorobiales as well as the HGT donor to a subset of Aquificales, a frequency observed HGT partner with Epsilonproteobacteria. However, long branches and sparse taxonomic representation prevent a reliable rooting and clear identification of a donor clade within Epsilonproteobacteria. The absence of aconitate hydratase within other photoheterotrophic lineages can be explained by two possible scenarios: (1) initial acquisition in the common ancestor lineage of *C. thalassium* and the photoheterotroph clade, with subsequent loss in the photoheterotroph ancestor; or (2) acquisition by *C. thalassium* after the divergence of the photoheterotroph ancestor lineage. Both scenarios include a subsequent independent acquisition by *Thermochlorobacter*. In the case of (1) the photoheterotroph clade can be inferred to be ancestrally autotrophic, as all rTCA cycle genes would be present in their common ancestor; in the case of (2) their common ancestor would be inferred to be lacking a complete rTCA cycle, and therefore extant photoheterotrophic lineages would represent a continuation of an ancestral metabolic state. However, this latter scenario is less likely, as all rTCA enzymes but aconitate hydratase would be ancestrally present, a “nearly complete” rather than merely a “partial” rTCA cycle that would not have a known modern analog.

**Figure 8.**
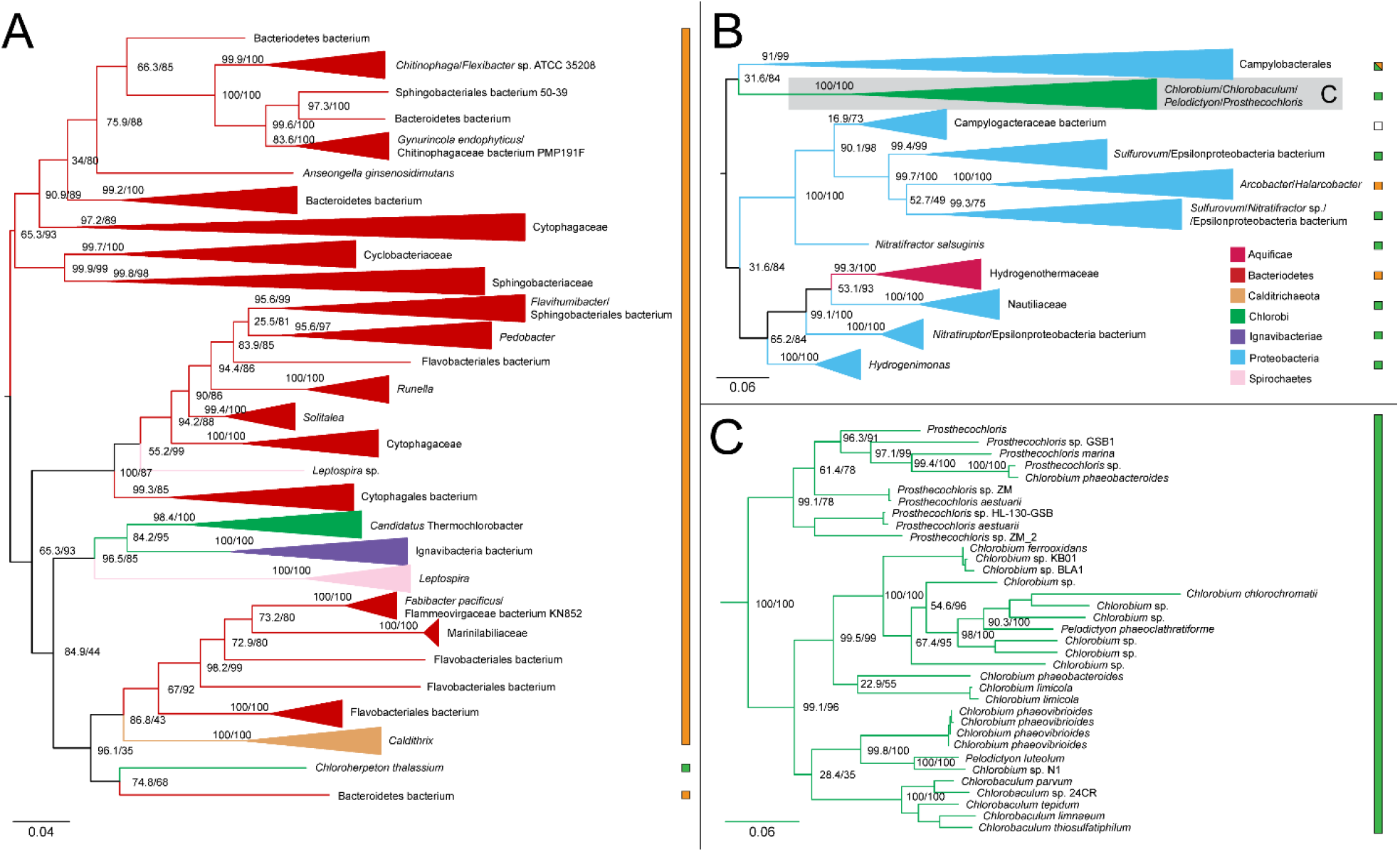
Maximum likelihood (ML) tree of aconitate hydratase homologs. Two groups of sequences related to Chlorobiales query sequences were identified, shown as midpoint rooted trees with collapsed clades labeled with taxonomic group names (A-B). A higher resolution tree for Chlorobiales sequences in tree B is shown in (C). Support values indicate approximate likelihood ratio test (aLRT)/bootstrap (100 replicates). Major bipartitions with bootstrap (BS) support are labeled. A and B trees contain 249 and 250 taxa, respectively. Color bars to the right of the tree indicate autotrophic (green), heterotrophic (orange), or undetermined (white) carbon metabolisms. Clades including multiple carbon metabolisms are indicated with diagonally hatched boxes.

### ATP Citrate Lyase

More than other enzymes, ATP citrate lyase is considered diagnostic for the presence of rTCA cycle carbon fixation, as it plays the crucial step of citrate cleavage into acetyl-CoA and oxaloacetate^38^. A variant of this process is found in Aquificae, where cleavage is catalyzed in tandem by citryl-CoA synthetase and citryl-CoA lyase^39^. These enzymes share a distant common ancestry indicated by sequence homology as well as conserved structural elements to the two subunits of ATP citrate lyase in *Chlorobium limicola*, specifically, a two-helix stalk and β-hairpins^40^ (Figure S2).

In our results, all autotrophic Chlorobiales form a single group in the ML phylogeny of both the α (Figure 9) and β (Figure S1) subunits of this protein, which includes Nitrospirae and Nitrospinae sequences, sibling to the deeply branching heterotrophic *Chlorobium* sp. 445 lineages (Figure 9B). The direction of HGT between Chlorobiales and Nitrospirae/Nitrospinae is unclear, as the recovered root of Chlorobiales is inconsistent with the species tree and varies between subunits. However, the α subunit sequence within *Chlorobium* sp. 445 is only partial, missing 167 amino acids (Figure S2). This absence in the alignment may impose tree reconstruction artifacts, resulting in the spurious rooting of this group, complicating HGT inference. To test this, we re-ran the ML tree using only the sequence region present in *Chlorobium* sp. 445. This recovered the same phylogenetic placement of sibling groups Nitrospirae/Nitrospinae with respect to Chlorobiales. Furthermore, Bayesian inference, an approach more resistant to long branch attraction artifacts, also recovered the same rooting for both subunits (Figure S3).

**Figure 9.**
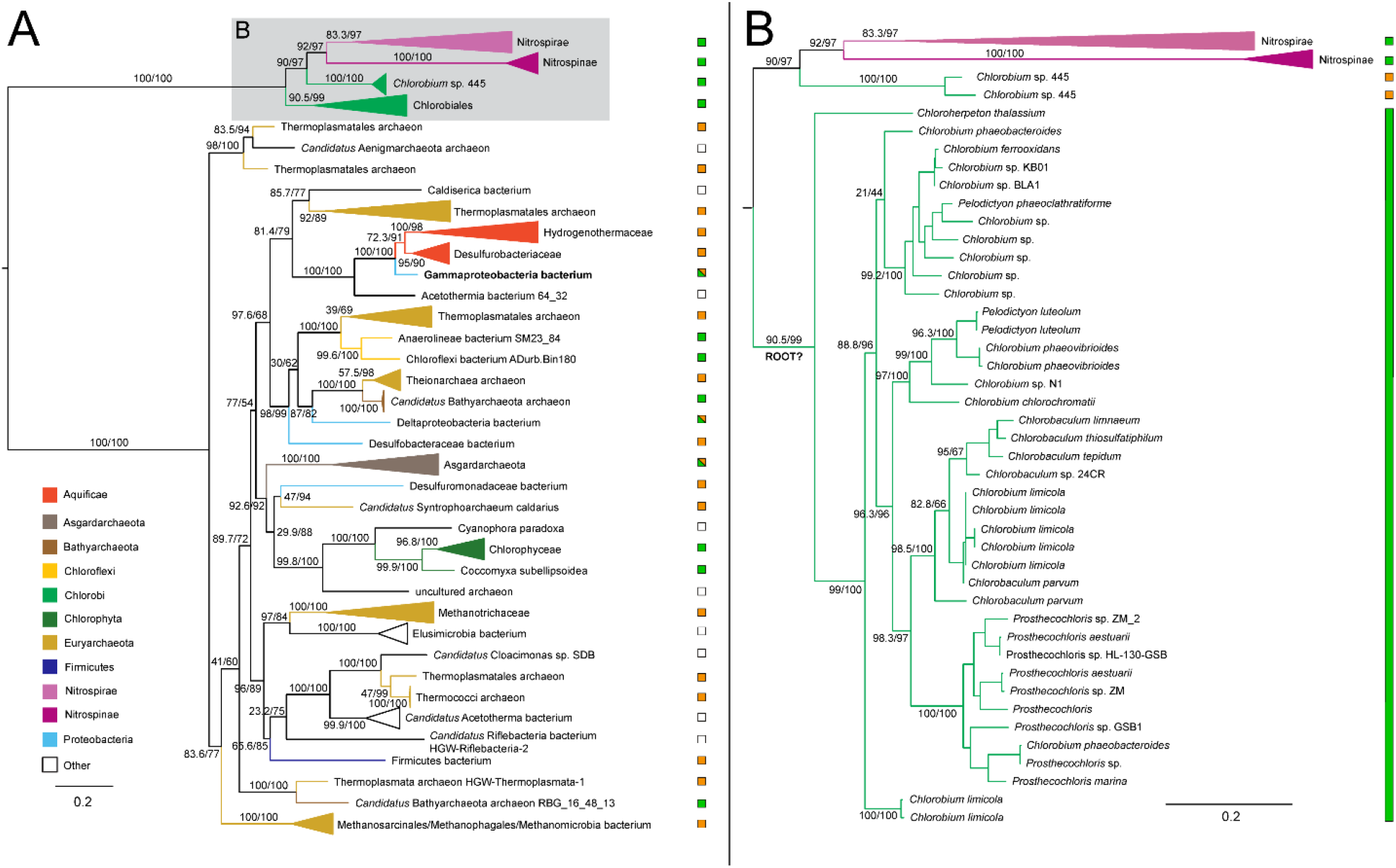
Maximum likelihood (ML) tree of ATP citrate lyase alpha subunit homologs. (A) Midpoint-rooted tree with collapsed clades labeled with taxonomic group names. (B) Higher resolution tree showing crown Chlorobiales and closely related sequences in Nitrospira/Nitrospinae. Support values indicate approximate likelihood ratio test (aLRT)/ bootstrap (100 replicates). Major clades with bootstrap (BS) support are labeled with respective values. The full tree contains 247 taxa. Color bars to the right of the tree indicate autotrophic (green), heterotrophic (orange), or undetermined (white) carbon metabolisms. Clades including multiple carbon metabolisms are indicated with diagonally hatched boxes.

A mapping of this missing region to the crystal structure of the ATP citrate lyase enzyme structure shows a missing a stabilizing β hairpin region (Figure S2), suggesting that this partial version may be nonfunctional, or retain a function unrelated to carbon fixation. This may also be evidence of independent loss of rTCA carbon fixation capability within other photoheterotrophic lineages, such as *Thermochlorobacter*. Still, without knowing if the truncated variant is functional, it brings the viability of ATP citrate lyase as a marker for autotrophy into question.

Other published phylogenies of ATP citrate lyase shows a history consistent with our findings, with vertical inheritance in Chlorobiales, but concluding that Nitrospirae and Nitrospinae horizontally transferred these genes to Chlorobiales, albeit with low statistical support^41^. The long branch separating these sequences from other taxa in the tree obscures the ancestry of ATP citrate lyase within the Chlorobiales stem lineage, and no clear HGT donor group can be inferred. The absence of other closely related sequences suggests an acquisition from unsampled or extinct groups. Further analyses are thus required to resolve the true history of these genes.

### Pyruvate: ferredoxin oxidoreductase

The ML tree for pyruvate: ferredoxin oxidoreductase shows very sparse taxonomic outgroups separated by a long branch indicative of independent HGT events. Distantly related groups include Bacteroidetes, Ignavibacteria, autotrophic Cyanobacteria, Chloroflexi, Verrucomicrobia, and Acidobacteria (Figure 10A). This enzyme is one of only three that are present in the rTCA cycle and not the oxidative TCA cycle^24^ and was apparently lost within the photoheterotrophic Chlorobiaceae lineages. Due to branch lengths, a putative donor group cannot be established.

**Figure 10.**
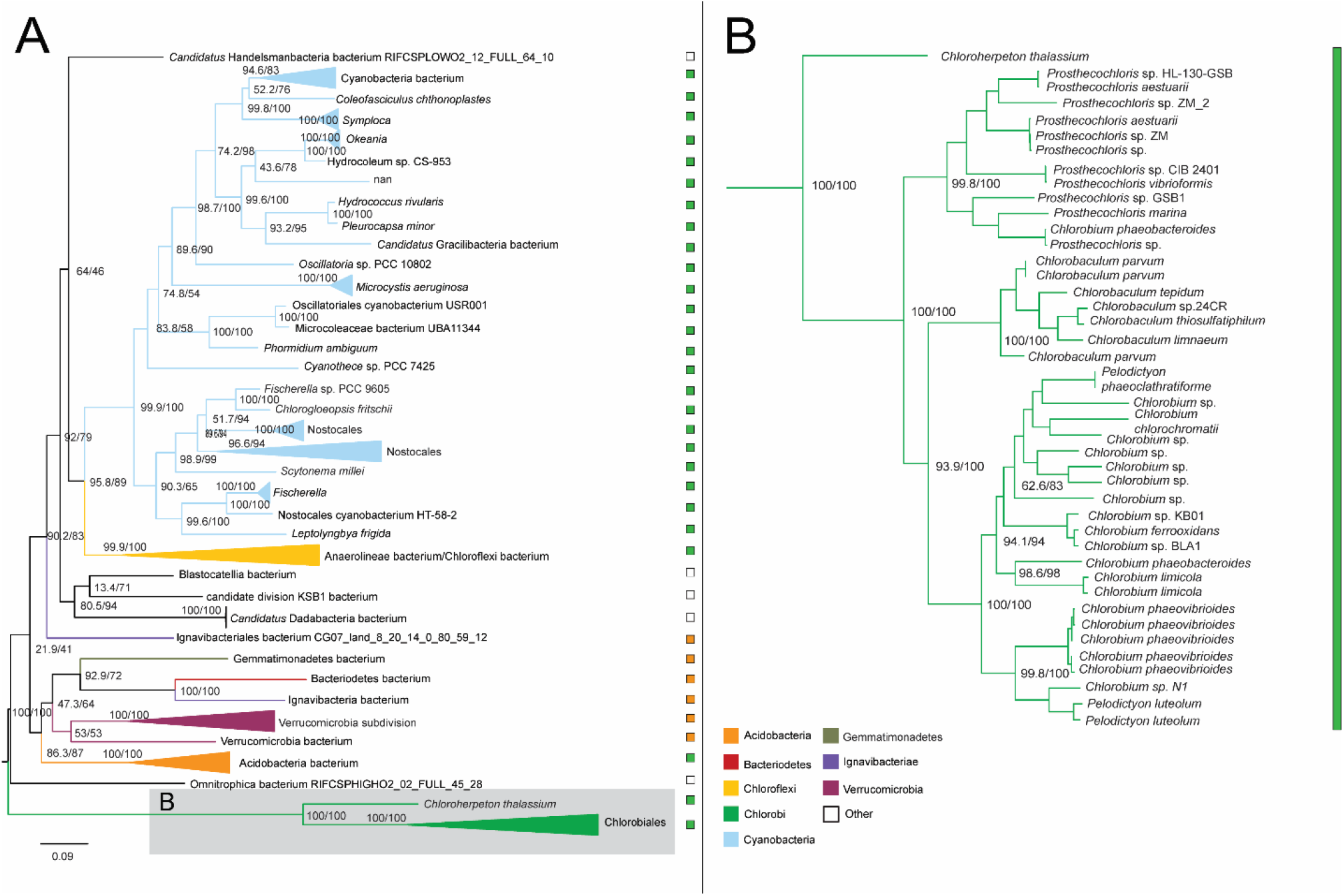
Maximum likelihood (ML) tree of pyruvate:ferredoxin oxidoreductase homologs. (A) Midpoint-rooted tree with collapsed clades labeled with taxonomic group names. (B) Higher resolution tree showing crown Chlorobiales and other closely related sequences. Support values indicate approximate likelihood ratio test (aLRT)/ bootstrap (100 replicates). Major clades with bootstrap (BS) support are labeled with respective values. The full tree contains 247 taxa. Color bars to the right of the tree indicate autotrophic (green), heterotrophic (orange), or undetermined (white) carbon metabolisms. Clades including multiple carbon metabolisms are indicated with diagonally hatched boxes.

### rTCA cycle proteins do not trace the evolutionary history of autotrophy

Additional evidence for the patchwork assembly of the rTCA cycle within Chlorobiales is provided by the taxonomic distribution of autotrophy across species represented in each protein tree. Surprisingly, in the majority of cases the most closely related outgroup sequences are not from autotrophic species, suggesting that these proteins have an alternative function in these groups (fumarate hydratase, succinyl-CoA synthetase, OGOR). Presumably, before the complete rTCA pathway was present, these proteins performed similar functions in Chlorobiales as part of a photoheterotrophic metabolism.

Further evidence of the potential utility of incomplete sets of rTCA cycle proteins is provided by the retention of several of these proteins within photoheterotrophic Chlorobiales; in some cases, it appears these proteins were acquired after the divergence from other Chlorobiaceae (fumarate hydratase, isocitrate lysase, aconitate hydratase) while others were vertically inherited within the crown group (malate dehydrogenase, succinyl-CoA synthetase 2-oxoglutarate: ferredoxin oxidoreductase). In all these cases, the genes present are also utilized in the oxidative TCA cycle, thus potentially explaining their retention in the divergent thermophiles. Elucidating their functions within these groups may provide clues to the ancestral functions of these genes in stem Chlorobiales, before a complete rTCA cycle was present. The inferred presence of ATP citrate lyase in stem Chlorobiales, in the absence of a complete and functional rTCA cycle, further suggests that this enzyme is not, on its own, diagnostic for autotrophy in this group, as is observed for other heterotrophic microbes^42,43^.

## Discussion

### A Chimeric origin of rTCA cycle genes and autotrophy within Chlorobiales

Our results show a striking diversity of the evolutionary histories of rTCA cycle enzymes within Chlorobiales, suggesting that carbon fixation has a complex and relatively recent evolutionary history within this group (Figure 11). There is strong evidence for the independent acquisition of several enzymes via HGT from different donors, both to the stem group, and to different lineages within Chlorobiales following their crown divergence. This interpretation is consistent with previous phylogenetic trees in which the inheritance of rTCA enzymes could not be narrowed down to a specific donor group or transfer event^41,44^.

**Figure 11.**
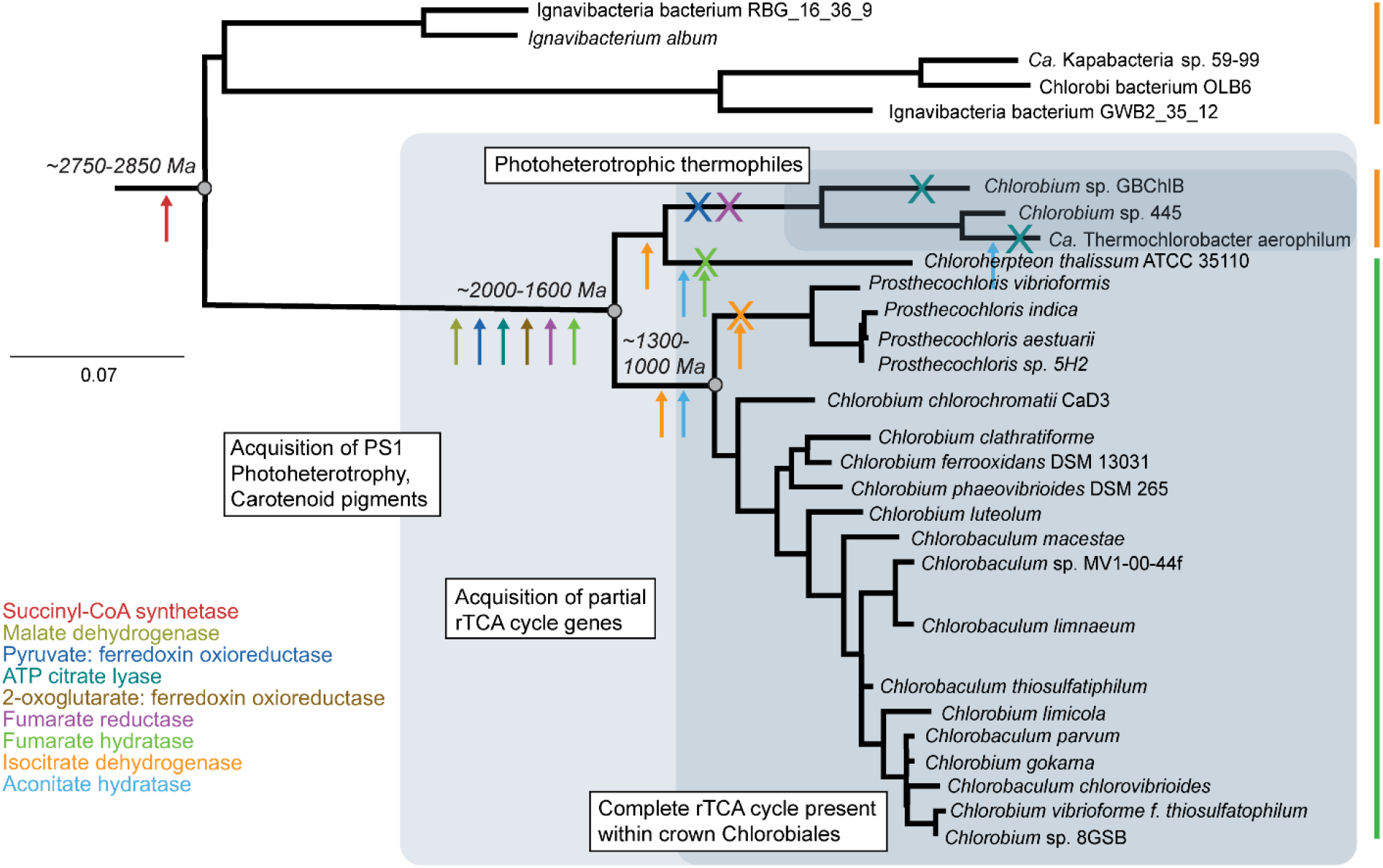
16S rRNA tree of Chlorobiales and Ignavibacterium (outgroup) with gene histories of rTCA proteins mapped. Arrows indicate inferred acquisitions of specific enzymes, and subsequent losses are indicated with crosses with enzyme names coded by color. Shaded fields indicate inferred carbon metabolisms for different clades. Divergence times are from Fournier et al., 2021 and are reported in SI data.

In several cases, *C. thalassium* and thermophilic GSB have orthologs of rTCA genes not closely related to those in other members of Chlorobiales, clearly indicating a more recent history of HGT for one or more subgroups. From these observations, two potential hypotheses emerge regarding the evolution of the rTCA cycle in Chlorobiales: (1) the full rTCA cycle was acquired in the stem lineage, with some enzymes replaced via additional HGTs in descendant crown group lineages; or (2) the full rTCA cycle was only assembled later, within crown Chlorobiales, with independent acquisition of some enzymes by multiple lineages. We find the latter explanation to be more parsimonious. Only for the fumarate hydratase tree is (1) a reasonable hypothesis (Fig. 3A). When *Candidatus Thermochlorobacter* and *Chlorobium sp. 445* appear with *C. thalassium* in the isocitrate lysase (Fig. 7A) and aconitate hydrase (Fig. 8A) trees, this implies (2) an independent acquisition via two HGT events following the divergence within the family of Chlorobi. These trees each have variable outgroups which include both autotrophic and heterotrophic bacteria/archaea.

### rTCA cycle genes within photoheterotrophic lineages

A functional rTCA cycle was likely originally present in the common ancestor of photoheterotrophic thermophilic GSB lineages. In several gene trees showing HGT within crown Chlorobiales, these group together with *C. thalassium*, indicating a likely shared ancestral state of autotrophy. Other genes present in a complete rTCA path were apparently lost in thermophilic GSB, consistent with their photoheterotrophic physiology. For example, *Candidatus Thermochlorobacter aerophilum* and *Chlorobium sp*. GBChlB do not have genes encoding ATP citrate lyase, an essential enzyme in the pathway (Figure 9B). Alternative genes to ATP citrate lyase that may rescue rTCA cycle function (citryl-CoA synthetase & citryl-CoA lyase)^39^ are also absent. The retention of a partial ATP citrate lyase within *Chlorobium*. sp. 455 further suggests that these genes were independently lost within thermophilic lineages following their diversification.

Other genes of the rTCA cycle were apparently retained in this group, suggesting functional roles independent of carbon fixation. It is also possible that 2-oxoglutarate:ferredoxin and pyruvate:ferredoxin might be involved in mixotrophic growth in an incomplete rTCA path in these lineages^33,34,45^.

### Molecular clocks inform the history of carbon fixation within Chlorobiales

The existence of the rTCA cycle as a primordial carbon fixation pathway, mediated by non-enzymatic, inorganic catalysts, is a hotly debated issue in origins of life research^20,46,47^. It has also been proposed that an ancestral rTCA was present in an autotrophic last universal common ancestor (LUCA)^48,49^. However, the phylogenomic signal needed to establish this deep ancestry may be impossible to fully elucidate, given the relatively shallow taxonomic distributions of rTCA cycle constituent proteins, and high frequency of HGT between phyla, as evidenced by reconstructed gene trees. Nevertheless, the clear history of rTCA cycle proteins within Chlorobiales, and the fidelity of their inheritance in several subgroups, permits the potential dating of carbon fixation within this clade. Recent molecular clock studies have produced age estimates for major divergences within Chlorobiales informed by large protein sequence datasets, cyanobacterial fossil calibrations, and HGT constraints^13^. These ages show crown Chlorobiales diversifying between 1977 Ma and 1607 Ma, and subsequent diversification of non-*Chloroherpeton* autotrophic lineages between 1300 Ma and 988 Ma (Figure 11). While these age uncertainties are large and will likely be revised in future studies with additional constraints, they are consistent with BCF lipid biomarkers at ∼1.64 Ga most likely being produced after the crown diversification of Chlorobiales but before the diversification of major extant autotrophic groups. These phylogenetic analyses therefore raise the intriguing possibility that BCF lipid biomarkers were produced by early diverging lineages that did not yet have a complete rTCA cycle. If that is indeed the case, they would not preserve C isotopic fractionations consistent with Chlorobiaceae carbon fixation under rTCA (Δδ^13^C -20 to -10%)^50^, but rather should show fractionations more like those observed for extant photoheterotrophic lineages, and/or BCF bulk organic carbon (Table 1). Furthermore, an isotopic shift in preserved GSB biomarkers to modern values reflective of carbon fixation via an rTCA cycle should be expected to occur before 988 Ma^13^. These interpretations may be complicated by the existence of unsampled or extinct Chlorobiales stem groups that preserved an ancestral heterotrophic metabolism long after the emergence of autotrophy within more derived groups; however, it is unlikely that such groups would be ecologically abundant following selection favoring autotrophy in these environments.

**Table 1:**
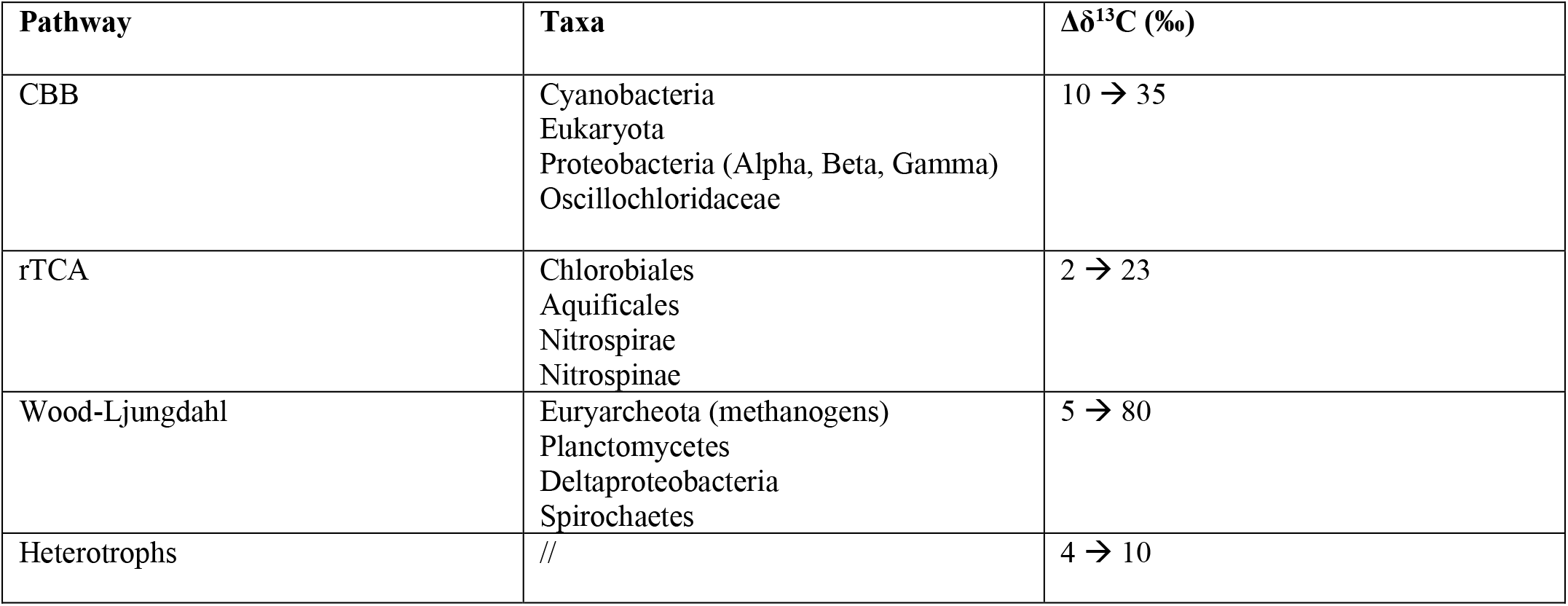
Isotopic differences by carbon fixation pathways and taxa^51–61^. Sourced as Δδ^13^C = δ^13^C_biomass_ −δ^13^C_carbon source_ calculation. Listed taxa aren’t diagnostic of full pathway distribution, but major groups of relevance. There is a ∼25‰ mean deviation between inorganic and organic carbon isotopes^62^.

### Autotrophy in Chlorobiales: adaptation to a changing Mesoproterozoic world?

The independent assembly of a complete rTCA cycle for carbon fixation in at least two major lineages of Chlorobiales is potentially indicative of broad changes in environmental conditions and ecological niches during the Mesoproterozoic. Because of their unique metabolisms and chemical requirements, GSB are useful indicators of past environmental conditions^63^, most notably for the changes in sulfur abundance and redox state through Earth history. The Mesoproterozoic was a time of complex interactions between sulfur and oxygen cycles, potentially spurring the metabolic evolution of anoxygenic phototrophs such as Chromaticeae and Chlorobiales. Redox-sensitive element abundances and stable isotopes indicate progressive oxygenation of the ocean–atmosphere system, along with a significant increase in marine sulfate inventory during this time^1,2,64–66^. This increased oxygenation of the atmosphere promoted weathering of pyrite and delivery of sulfate to aquatic environments, subsequently increasing the prevalence of euxinic conditions via microbial sulfate reduction^67,68^. At the same time, increased oxygenation of surface waters would drive anoxygenic phototrophs deeper in water columns where these anoxic conditions could persist; these lower light environments would select for more efficient light-harvesting systems, such as chlorosomes^69^, and more energy efficient metabolic pathways for processing carbon. This may also explain why Chlorobiales and Chromatiales don’t have shared carbon fixation pathways despite their similar physiology and environmental occurrence. Chromatiales are more oxygen-tolerant and can exist higher in the water column where light is more available for energy-intensive carbon fixation via the CBB cycle, requiring seven ATP per molecule of synthesized pyruvate, compared with only two for the rTCA cycle^38^.

In such a scenario, the HGT donors of rTCA cycle genes to Chlorobiales would also have inhabited these deeper aquatic environments. Unfortunately, the phylogenies of rTCA cycle proteins do not identify putative HGT donor groups with sufficient resolution to evaluate this hypothesis with the present dataset. Future environmental sequencing investigations of stratified marine environments may discover missing microbial diversity that better resolves these gene tree histories; alternatively, these microbial groups and their metabolisms may have been unique to stratified Proterozoic marine conditions, and have since gone extinct, or have lost these genes. While this scenario may explain why Chlorobiales acquired carbon fixation via rTCA cycle genes rather than CBB cycle genes, it does not, on its own, explain why the shift to autotrophy occurred during this time. One possibility may once again relate to increasing oxygen levels. If aquatic aerobic/sulfate reducing metabolisms also experienced a diversification and ecological expansion at this time, then, presumably, more organic carbon would be microbially oxidized, leaving less available for assimilatory heterotrophic processes^70^. These conditions would therefore favor more widespread autotrophy, even among energy-limited organisms such as Chlorobiales.

## Conclusion

Our proposed scenario for the history of autotrophy within Chlorobiales infers early members of this group were heterotrophs and that autotrophy was independently acquired within crown Chlorobiales lineages through a chimeric history of acquisition of rTCA genes. Therefore, early members of Chlorobiales fractionated carbon differently than extant autotrophic members that have a complete rTCA cycle, a transition that is predicted to be apparent within the isotopic record of lipid biomarkers. Molecular clock studies of Chlorobiales constrain the timing of these metabolic evolutionary events, providing the means to integrate these genomic and geochemical records to establish a well-resolved evolutionary ecology of GSB. This work demonstrates how phylogenomic analysis can provide an independent source of information for interpreting the lipid biomarker record, and the importance of integrating phylogenomic data with stable carbon isotope analysis in inferring the evolutionary history of microbial metabolisms.

## Methods

### Genomic Data Retrieval

Query sequences for rTCA cycle enzymes were selected from the sequenced *Chlorobium thalissium ATCC 35110* genome (CP001100.1) retrieved from the NCBI database (SI Table 1). In cases where the protein ortholog from *Chloroherpeton thalissum* did not recover monophyly with other Chlorobiaceae, orthologous query sequences were taken from *Chlorobium tepidum TLS* (AE006470.1).

### BLAST Search for Homologous Enzymes

The Basic Local Alignment Search Tool (BLAST) was used to compare queries with homologous sequences using default search parameters. The BLASTp cutoff for homologous identification was set at 250 taxa maximum, with sequence identity ≥ 32%. These searchers were not exhaustive for all rTCA homologs, but rather exhaustively identified all Chlorobiales homologs and extensively sampled sequences representing potential HGT donor lineages and/or distant vertically inherited sequences from outgroup clades.

### Phylogenetic Analysis

Protein sequences were aligned using the multiple sequence alignment tool MAFFT v.7.245 (FFT-NS-2, BLOSUM62)^71^. Maximum-likelihood phylogenetic trees were generated using IQTree run with the Bayes Information Criterion (BIC) test (SI Table 1). Support for bipartitions was inferred using rapid bootstraps (1000 replicates) and SH-aLRT tests. Additional phylogenetic analysis of the ATP citrate lyase dataset was performed using Phylobayes v. 4.1^71^, using C20 site specific profiles and a convergence criterion cutoff of 0.3 for all parameters, with a 20% chain burn-in. Trees were rooted at midpoints and visualized in FigTree v.1.4.4.

### Data Availability

Supplementary data files are available at Dryad DOI: https://doi.org/10.5061/dryad.931zcrjmw

## Notes

https://doi.org/10.5061/dryad.931zcrjmw

### Competing Interest Statement

The authors have declared no competing interest.

## References

1. Lyons TW, Reinhard CT, Planavsky NJ. The rise of oxygen in Earth’s early ocean and atmosphere. Nature. 2014;506(7488):307–315. doi:10.1038/nature13068

2. Holland HD. The oxygenation of the atmosphere and oceans. Philos Trans R Soc B Biol Sci. 2006;361(1470):903–915. doi:10.1098/rstb.2006.1838

3. Briggs DEG, Summons RE. Ancient biomolecules: their origins, fossilization, and role in revealing the history of life. BioEssays News Rev Mol Cell Dev Biol. 2014;36(5):482–490. doi:10.1002/bies.201400010

4. Isorenieratane record in black shales from the Paris Basin, France: Constraints on recycling of respired CO2 as a mechanism for negative carbon isotope shifts during the Toarcian oceanic anoxic event - van Breugel - 2006 - Paleoceanography - Wiley Online Library. Accessed January 20, 2022. https://agupubs.onlinelibrary.wiley.com/doi/full/10.1029/2006PA001305

5. Brocks JJ, Schaeffer P. Okenane, a biomarker for purple sulfur bacteria (Chromatiaceae), and other new carotenoid derivatives from the 1640Ma Barney Creek Formation. Geochim Cosmochim Acta. 2008;72(5):1396–1414. doi:10.1016/j.gca.2007.12.006

6. Brocks JJ, Love GD, Summons RE, Knoll AH, Logan GA, Bowden SA. Biomarker evidence for green and purple sulphur bacteria in a stratified Palaeoproterozoic sea. Nature. 2005;437(7060):866–870. doi:10.1038/nature04068

7. Reinhard CT, Planavsky NJ, Robbins LJ, et al. Proterozoic ocean redox and biogeochemical stasis. Proc Natl Acad Sci. 2013;110(14):5357–5362. doi:10.1073/pnas.1208622110

8. Klassen JL. Phylogenetic and Evolutionary Patterns in Microbial Carotenoid Biosynthesis Are Revealed by Comparative Genomics. PLOS ONE. 2010;5(6):e11257. doi:10.1371/journal.pone.0011257

9. Maresca JA, Graham JE, Bryant DA. The biochemical basis for structural diversity in the carotenoids of chlorophototrophic bacteria. Photosynth Res. 2008;97(2):121–140. doi:10.1007/s11120-008-9312-3

10. Graham JE, Bryant DA. The Biosynthetic Pathway for Synechoxanthin, an Aromatic Carotenoid Synthesized by the Euryhaline, Unicellular Cyanobacterium Synechococcus sp. Strain PCC 7002. J Bacteriol. Published online December 2008. doi:10.1128/JB.00985-08

11. Cui X, Liu XL, Shen G, et al. Niche expansion for phototrophic sulfur bacteria at the Proterozoic–Phanerozoic transition. Proc Natl Acad Sci. 2020;117(30):17599–17606. doi:10.1073/pnas.2006379117

12. Magnabosco C, Moore KR, Wolfe JM, Fournier GP. Dating phototrophic microbial lineages with reticulate gene histories. Geobiology. 2018;16(2):179–189. doi:10.1111/gbi.12273

13. Fournier GP, Moore KR, Rangel LT, Payette JG, Momper L, Bosak T. The Archean origin of oxygenic photosynthesis and extant cyanobacterial lineages. Proc R Soc B Biol Sci. 2021;288(1959):20210675. doi:10.1098/rspb.2021.0675

14. Hayes JM. 3. Fractionation of Carbon and Hydrogen Isotopes in Biosynthetic Processes. In: 3. Fractionation of Carbon and Hydrogen Isotopes in Biosynthetic Processes. De Gruyter; 2018:225–278. doi:10.1515/9781501508745-006

15. Jennings R de M, Moran JJ, Jay ZJ, et al. Integration of Metagenomic and Stable Carbon Isotope Evidence Reveals the Extent and Mechanisms of Carbon Dioxide Fixation in High-Temperature Microbial Communities. Front Microbiol. 2017;8. Accessed February 4, 2022. https://www.frontiersin.org/article/10.3389/fmicb.2017.00088

16. Ward LM, Shih PM. The evolution and productivity of carbon fixation pathways in response to changes in oxygen concentration over geological time. Free Radic Biol Med. 2019;140:188–199. doi:10.1016/j.freeradbiomed.2019.01.049

17. Tabita FR, Hanson TE, Satagopan S, Witte BH, Kreel NE. Phylogenetic and evolutionary relationships of RubisCO and the RubisCO-like proteins and the functional lessons provided by diverse molecular forms. Philos Trans R Soc Lond B Biol Sci. 2008;363(1504):2629–2640. doi:10.1098/rstb.2008.0023

18. Hügler M, Huber H, Stetter KO, Fuchs G. Autotrophic CO2 fixation pathways in archaea (Crenarchaeota). Arch Microbiol. 2003;179(3):160–173. doi:10.1007/s00203-002-0512-5

19. Hügler M, Wirsen CO, Fuchs G, Taylor CD, Sievert SM. Evidence for Autotrophic CO2 Fixation via the Reductive Tricarboxylic Acid Cycle by Members of the ε Subdivision of Proteobacteria. J Bacteriol. Published online May 1, 2005. doi:10.1128/JB.187.9.3020-3027.2005

20. Wächtershäuser G. Evolution of the first metabolic cycles. Proc Natl Acad Sci. 1990;87(1):200–204. doi:10.1073/pnas.87.1.200

21. Braakman R, Smith E. The Emergence and Early Evolution of Biological Carbon-Fixation. PLOS Comput Biol. 2012;8(4):e1002455. doi:10.1371/journal.pcbi.1002455

22. Srinivasan V, Morowitz HJ. The canonical network of autotrophic intermediary metabolism: minimal metabolome of a reductive chemoautotroph. Biol Bull. 2009;216(2):126–130. doi:10.1086/BBLv216n2p126

23. Ivanovsky RN, Sintsov NV, Kondratieva EN. ATP-linked citrate lyase activity in the green sulfur bacterium Chlorobium limicola forma thiosulfatophilum. Arch Microbiol. Published online 2004. doi:10.1007/BF00406165

24. Tang KH, Blankenship RE. Both Forward and Reverse TCA Cycles Operate in Green Sulfur Bacteria. J Biol Chem. 2010;285(46):35848–35854. doi:10.1074/jbc.M110.157834

25. Nakano MM, Zuber P, Sonenshein AL. Anaerobic regulation of Bacillus subtilis Krebs cycle genes. J Bacteriol. 1998;180(13):3304–3311. doi:10.1128/JB.180.13.3304-3311.1998

26. Aoshima M. Novel enzyme reactions related to the tricarboxylic acid cycle: phylogenetic/functional implications and biotechnological applications. Appl Microbiol Biotechnol. 2007;75(2):249–255. doi:10.1007/s00253-007-0893-0

27. Hügler M, Huber H, Molyneaux SJ, Vetriani C, Sievert SM. Autotrophic CO2 fixation via the reductive tricarboxylic acid cycle in different lineages within the phylum Aquificae: evidence for two ways of citrate cleavage. Environ Microbiol. 2007;9(1):81–92. doi:10.1111/j.1462-2920.2006.01118.x

28. Gunsalus RP, Park SJ. Aerobic-anaerobic gene regulation in Escherichia coli: control by the ArcAB and Fnr regulons. Res Microbiol. 1994;145(5-6):437–450. doi:10.1016/0923-2508(94)90092-2

29. Ma K, Hutchins A, Sung SJ, Adams MW. Pyruvate ferredoxin oxidoreductase from the hyperthermophilic archaeon, Pyrococcus furiosus, functions as a CoA-dependent pyruvate decarboxylase. Proc Natl Acad Sci U S A. 1997;94(18):9608–9613. doi:10.1073/pnas.94.18.9608

30. Imhoff JF. Phylogenetic taxonomy of the family Chlorobiaceae on the basis of 16S rRNA and fmo (Fenna-Matthews-Olson protein) gene sequences. Int J Syst Evol Microbiol. 2003;53(Pt 4):941–951. doi:10.1099/ijs.0.02403-0

31. Olsen GJ, Woese CR, Overbeek R. The winds of (evolutionary) change: breathing new life into microbiology. J Bacteriol. 1994;176(1):1–6.

32. Hiras J, Wu YW, Eichorst SA, Simmons BA, Singer SW. Refining the phylum Chlorobi by resolving the phylogeny and metabolic potential of the representative of a deeply branching, uncultivated lineage. ISME J. 2016;10(4):833–845. doi:10.1038/ismej.2015.158

33. Klatt CG, Wood JM, Rusch DB, et al. Community ecology of hot spring cyanobacterial mats: predominant populations and their functional potential. ISME J. 2011;5(8):1262–1278. doi:10.1038/ismej.2011.73

34. Liu Z, Klatt CG, Ludwig M, et al. ‘Candidatus Thermochlorobacter aerophilum:’ an aerobic chlorophotoheterotrophic member of the phylum Chlorobi defined by metagenomics and metatranscriptomics. ISME J. 2012;6(10):1869–1882. doi:10.1038/ismej.2012.24

35. Stamps BW, Corsetti FA, Spear JR, Stevenson BS. Draft Genome of a Novel Chlorobi Member Assembled by Tetranucleotide Binning of a Hot Spring Metagenome. Genome Announc. Published online September 11, 2014. doi:10.1128/genomeA.00897-14

36. Speth DR, in ’t Zandt MH, Guerrero-Cruz S, Dutilh BE, Jetten MSM. Genome-based microbial ecology of anammox granules in a full-scale wastewater treatment system. Nat Commun. 2016;7(1):11172. doi:10.1038/ncomms11172

37. Phillips D, Aponte AM, French SA, Chess DJ, Balaban RS. Succinyl-CoA synthetase is a phosphate target for the activation of mitochondrial metabolism. Biochemistry. 2009;48(30):7140–7149. doi:10.1021/bi900725c

38. Hügler M, Sievert SM. Beyond the Calvin Cycle: Autotrophic Carbon Fixation in the Ocean. Annu Rev Mar Sci. 2011;3(1):261–289. doi:10.1146/annurev-marine-120709-142712

39. Aoshima M, Ishii M, Igarashi Y. A novel enzyme, citryl-CoA synthetase, catalysing the first step of the citrate cleavage reaction in Hydrogenobacter thermophilus TK-6. Mol Microbiol. 2004;52(3):751–761. doi:10.1111/j.1365-2958.2004.04009.x

40. Verschueren KHG, Blanchet C, Felix J, et al. Structure of ATP citrate lyase and the origin of citrate synthase in the Krebs cycle. Nature. 2019;568(7753):571–575. doi:10.1038/s41586-019-1095-5

41. Ward L, Shih P. Phototrophy and Carbon Fixation in Chlorobi Postdate the Rise of Oxygen.; 2021. doi:10.1101/2021.01.22.427768

42. Möller D, Schauder R, Fuchs G, Thauer RK. Acetate oxidation to CO2 via a citric acid cycle involving an ATP-citrate lyase: a mechanism for the synthesis of ATP via substrate level phosphorylation in Desulfobacter postgatei growing on acetate and sulfate. Arch Microbiol. 1987;148(3):202–207. doi:10.1007/BF00414812

43. Schauder R, Widdel F, Fuchs G. Carbon assimilation pathways in sulfate-reducing bacteria II. Enzymes of a reductive citric acid cycle in the autotrophic Desulfobacter hydrogenophilus. Arch Microbiol. Published online 2004. doi:10.1007/BF00414815

44. Becerra A, Rivas M, García-Ferris C, Lazcano A, Peretó J. A phylogenetic approach to the early evolution of autotrophy: the case of the reverse TCA and the reductive acetyl-CoA pathways. Int Microbiol Off J Span Soc Microbiol. 2014;17(2):91–97. doi:10.2436/20.1501.01.211

45. Bryant DA, Liu Z, Li T, et al. Comparative and Functional Genomics of Anoxygenic Green Bacteria from the Taxa Chlorobi, Chloroflexi, and Acidobacteria. In: Burnap R, Vermaas W, eds. Functional Genomics and Evolution of Photosynthetic Systems. Advances in Photosynthesis and Respiration. Springer Netherlands; 2012:47–102. doi:10.1007/978-94-007-1533-2_3

46. Smith E, Morowitz HJ. Universality in intermediary metabolism. Proc Natl Acad Sci. 2004;101(36):13168–13173. doi:10.1073/pnas.0404922101

47. Orgel LE. The Implausibility of Metabolic Cycles on the Prebiotic Earth. PLOS Biol. 2008;6(1):e18. doi:10.1371/journal.pbio.0060018

48. Wächtershäuser G. Before enzymes and templates: theory of surface metabolism. Microbiol Rev. 1988;52(4):452–484.

49. Weiss MC, Sousa FL, Mrnjavac N, et al. The physiology and habitat of the last universal common ancestor. Nat Microbiol. 2016;1(9):1–8. doi:10.1038/nmicrobiol.2016.116

50. Schidlowski M. Carbon isotopes as biogeochemical recorders of life over 3.8 Ga of Earth history: evolution of a concept. Precambrian Res. 2001;106(1):117–134. doi:10.1016/S0301-9268(00)00128-5

51. McNevin DB, Badger MR, Whitney SM, von Caemmerer S, Tcherkez GGB, Farquhar GD. Differences in Carbon Isotope Discrimination of Three Variants of D-Ribulose-1,5-bisphosphate Carboxylase/Oxygenase Reflect Differences in Their Catalytic Mechanisms*♦. J Biol Chem. 2007;282(49):36068–36076. doi:10.1074/jbc.M706274200

52. House CH, Schopf JW, Stetter KO. Carbon isotopic fractionation by Archaeans and other thermophilic prokaryotes. Org Geochem. 2003;34(3):345–356. doi:10.1016/S0146-6380(02)00237-1

53. Preuß A, Schauder R, Fuchs G, Stichler W. Carbon Isotope Fractionation by Autotrophic Bacteria with Three Different C02 Fixation Pathways. Z Für Naturforschung C. 1989;44(5-6):397–402. doi:10.1515/znc-1989-5-610

54. Quandt L, Gottschalk G, Ziegler H, Stichler W. Isotope discrimination by photosynthetic bacteria. FEMS Microbiol Lett. 1977;1(3):125–128. doi:10.1111/j.1574-6968.1977.tb00596.x

55. Zyakun AM, Lunina ON, Prusakova TS, Pimenov NV, Ivanov MV. Fractionation of stable carbon isotopes by photoautotrophically growing anoxygenic purple and green sulfur bacteria. Microbiology. 2009;78(6):757. doi:10.1134/S0026261709060137

56. Londry KL, Dawson KG, Grover HD, Summons RE, Bradley AS. Stable carbon isotope fractionation between substrates and products of Methanosarcina barkeri. Org Geochem. 2008;39(5):608–621. doi:10.1016/j.orggeochem.2008.03.002

57. Penger J, Conrad R, Blaser M. Stable Carbon Isotope Fractionation by Methylotrophic Methanogenic Archaea | Applied and Environmental Microbiology. Published 2012. Accessed February 4, 2022. https://journals.asm.org/doi/10.1128/AEM.01773-12

58. Gelwicks JT, Risatti JB, Hayes JM. Carbon isotope effects associated with autotrophic acetogenesis. Org Geochem. 1989;14(4):441–446. doi:10.1016/0146-6380(89)90009-0

59. Figueroa IA, Barnum TP, Somasekhar PY, Carlström CI, Engelbrektson AL, Coates JD. Metagenomics-guided analysis of microbial chemolithoautotrophic phosphite oxidation yields evidence of a seventh natural CO2 fixation pathway. Proc Natl Acad Sci. 2018;115(1):E92–E101. doi:10.1073/pnas.1715549114

60. Min K, Lehmeier C, Ballantyne IV F, Billings S. Frontiers | Carbon Availability Modifies Temperature Responses of Heterotrophic Microbial Respiration, Carbon Uptake Affinity, and Stable Carbon Isotope Discrimination | Microbiology. Published 2016. Accessed February 4, 2022. https://www.frontiersin.org/articles/10.3389/fmicb.2016.02083/full

61. Garcia AK, Cavanaugh CM, Kacar B. The curious consistency of carbon biosignatures over billions of years of Earth-life coevolution. ISME J. 2021;15(8):2183–2194. doi:10.1038/s41396-021-00971-5

62. Raven JA. Contributions of anoxygenic and oxygenic phototrophy and chemolithotrophy to carbon and oxygen fluxes in aquatic environments. Aquat Microb Ecol. 2009;56(2-3):177–192. doi:10.3354/ame01315

63. Menzel D, Hopmans EC, Bergen PF van, Leeuw JW de, Damsté JSS. Development of photic zone euxinia in the eastern Mediterranean Basin during deposition of Pliocene sapropels. Mar Geol. 2002;3-4(189):215–226. doi:10.1016/S0025-3227(02)00479-6

64. Canfield DE. The evolution of the Earth surface sulfur reservoir. Am J Sci. 2004;304(10):839–861. doi:10.2475/ajs.304.10.839

65. Halverson GP, Hurtgen MT. Ediacaran growth of the marine sulfate reservoir. Earth Planet Sci Lett. 2007;263(1-2):32–44. doi:10.1016/j.epsl.2007.08.022

66. Blättler CL, Bergmann KD, Kah LC, Gómez-Pérez I, Higgins JA. Constraints on Meso-to Neoproterozoic seawater from ancient evaporite deposits. Earth Planet Sci Lett. 2020;532:115951. doi:10.1016/j.epsl.2019.115951

67. Berner RA, Canfield DE. A new model for atmospheric oxygen over Phanerozoic time. Am J Sci. 1989;289(4):333–361. doi:10.2475/ajs.289.4.333

68. Hurtgen MT, Arthur MA, Halverson GP. Neoproterozoic sulfur isotopes, the evolution of microbial sulfur species, and the burial efficiency of sulfide as sedimentary pyrite. Geology. 2005;33(1):41–44. doi:10.1130/G20923.1

69. Ozaki K, Thompson K, Simister R, Crowe S, Reinhard CT. Anoxygenic photosynthesis and the delayed oxygenation of Earth’s atmosphere | Nature Communications. Published 2019. Accessed February 4, 2022. https://www.nature.com/articles/s41467-019-10872-z

70. Jørgensen BB, Findlay AJ, Pellerin A. Frontiers | The Biogeochemical Sulfur Cycle of Marine Sediments | Microbiology. Published 2019. Accessed February 4, 2022. https://www.frontiersin.org/articles/10.3389/fmicb.2019.00849/full

71. Katoh K, Standley DM. MAFFT Multiple Sequence Alignment Software Version 7: Improvements in Performance and Usability. Mol Biol Evol. 2013;30(4):772–780. doi:10.1093/molbev/mst010

