## Supplemental for "Chimeric Inheritance and Crown-Group Acquisitions of Carbon Fixation Genes within Chlorobiales"

SI Table 1. Enzymes, NCBI Accessions, Genome IDs, alignment, taxonomic diversity, and ML tree model for query sequences.

| **Enzyme Name** | **Accession** | **Genome ID** | **Alignment Length** | **# Taxa** | **BIC** |
|---|---|---|---|---|---|
| malate dehydrogenase | WP_012498653 | CP001100.1 | 318 | 242 | LG+R6 |
| fumarate hydratase | WP_012500396 | CP001100.1 | 480 | 250 | LG+R8 |
| fumarate hydratase | WP_010932514 | AE006470.1 | 565 | 242 | LG+R7 |
| fumarate reductase | WP_012498718 | CP001100.1 | 448 | 248 | LG+R8 |
| succinyl-CoA synthetase | WP_012500801 | CP001100.1 | 426 | 248 | LG+R7 |
| 2-oxoglutarate:ferredoxin oxidoreductase, alpha subunit | WP_012500035 | CP001100.1 | 657 | 243 | LG+R7 |
| isocitrate dehydrogenase | ACF14612 | CP001100.1 | 356 | 232 | LG+R8 |
| isocitrate dehydrogenase | WP_010932043 | AE006470.1 | 765 | 250 | LG+F+R7 |
| aconitate hydratase | WP_012500984 | CP001100.1 | 775 | 249 | LG+F+R8 |
| aconitate hydratase | WP_010932230 | AE006470.1 | 889 | 250 | LG+F+R7 |
| ATP citrate lyase alpha subunit | WP_012501083 | CP001100.1 | 658 | 247 | LG+R7 |
| ATP citrate lyase beta subunit | WP_012501083 | CP001100.1 | 443 | 287 | LG+R7 |
| pyruvate:ferredoxin oxidoreductase | WP_012499296 | CP001100.1 | 1357 | 247 | LG+R6 |

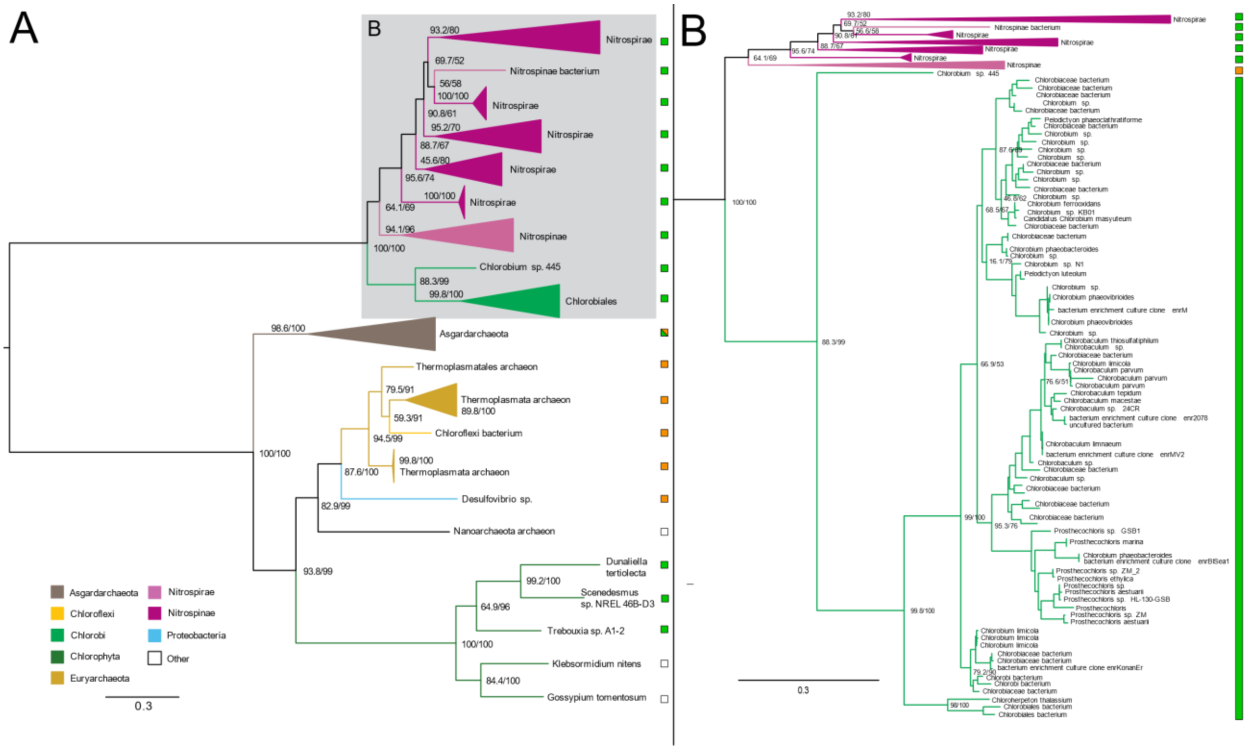


**SI Figure 1:** Maximum likelihood (ML) tree of ATP citrate lyase beta subunit homologs. (A) midpoint-rooted tree with collapsed clades labeled with taxonomic group names. (B) Higher resolution tree showing crown Chlorobiales and closely related sequences in Nitrospira/Nitrospinae. Support values indicate approximate likelihood ratio test (aLRT)/ bootstrap (100 replicates). Major clades with bootstrap (BS) support are labeled with respective values. Color bars to the right of the tree indicate autotrophic (green), heterotrophic (orange), or undetermined (white) carbon metabolisms.

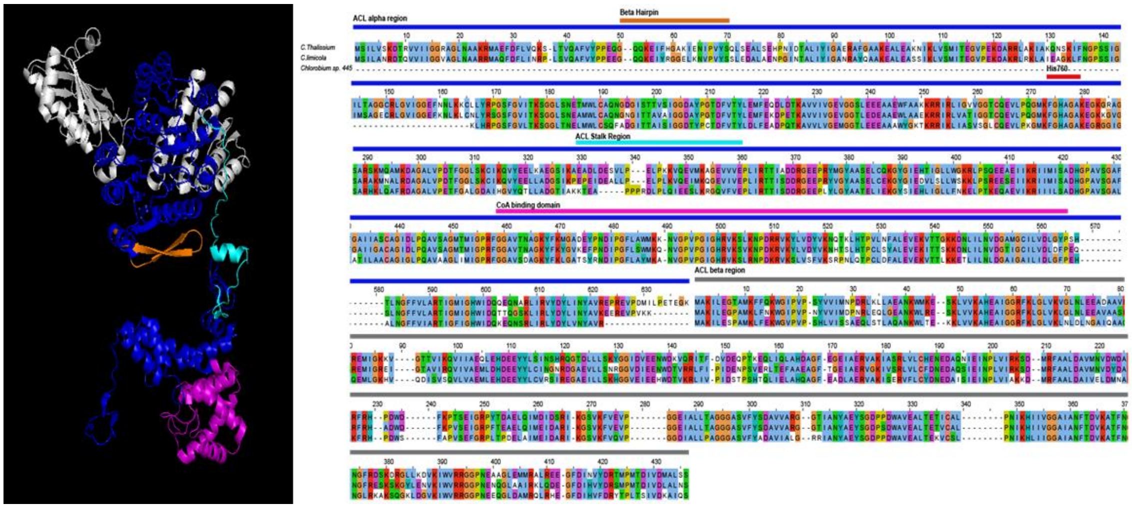


**SI Figure 2:** Protein Structural map of *Chlorobium Limicola* ATP Citrate Lyase enzyme (cite published structure here). Structural and labeled functional regions regions are colored corresponding to bars above sequence alignments. The amino acid alignment includes are sequences from *C.thalassium*, *C.limicola*, *Chlorobium* sp.445.

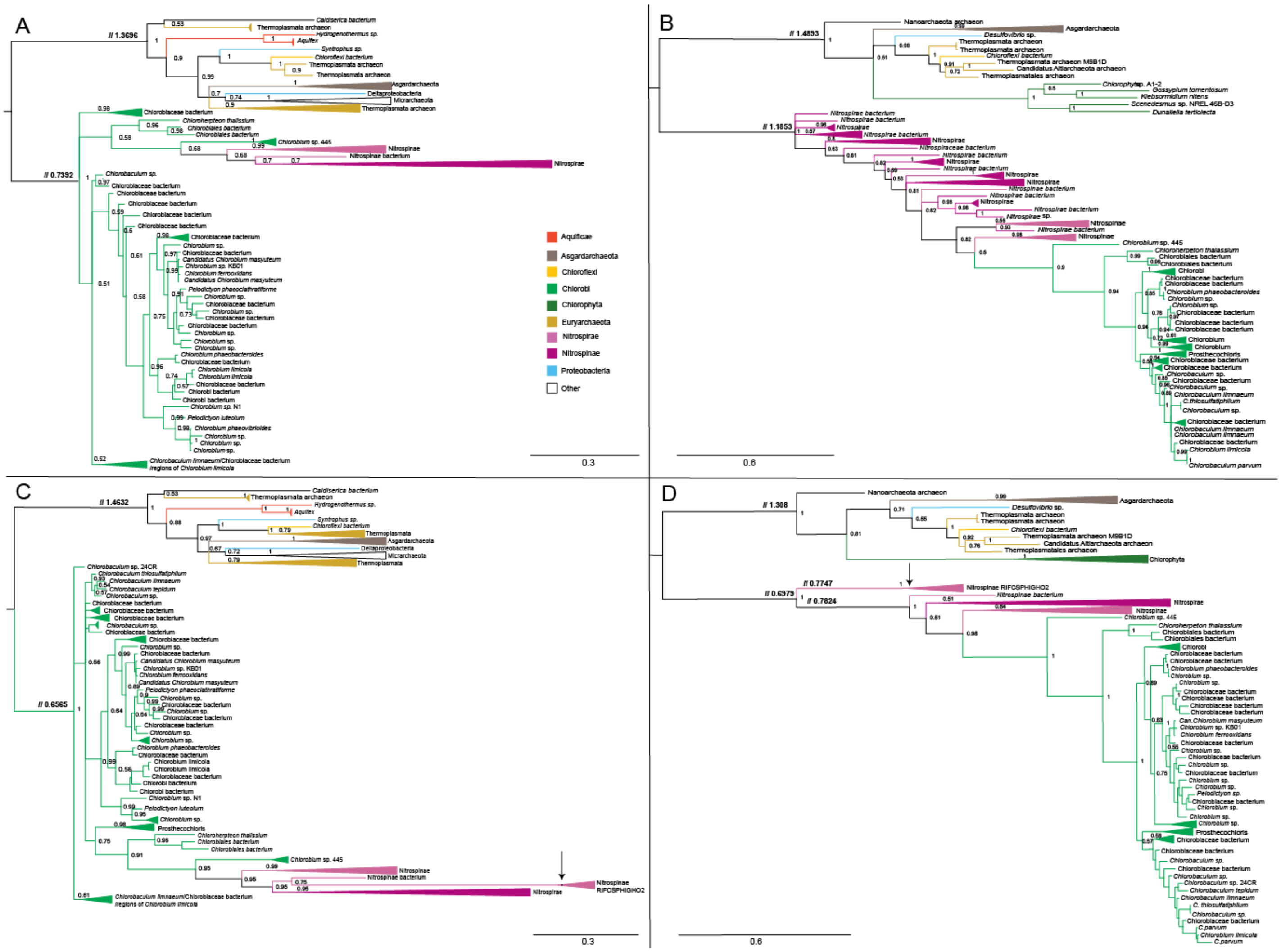


**SI Figure 3:** Bayesian Inference (BI) consensus trees of ATP citrate lyase alpha (A, C) and beta (B, D) subunit homologs. Trees depicted without (A, B) and with (C, D) Nitrospinae metagenomic sequences from He et. al., 2020 included in the alignment depicted with arrow. Collapsed clades labeled with taxonomic group names. Support values show consensus posterior probabilities.
